# PIP_2_-TMIE Interactions Drive Mammalian Hair Cell Slow Adaptation Independently of Myosin Motors

**DOI:** 10.1101/2025.04.01.646713

**Authors:** Giusy A. Caprara, Sujin Jun, Ye-Ri Kim, Gabriel J. Olguín-Orellana, Yein Christina Park, Claudia Martínez-García, Sihan Li, Angela Ballesteros, David Ramírez, Unkyung Kim, Jung-Bum Shin, Anthony W. Peng

## Abstract

Sensory hair cells detect sound and balance through their apically located stereocilia bundles, converting mechanical stimuli into electrical signals via mechano-electrical transduction (MET) channels. These channels at the lower end of extracellular tip links connecting adjacent stereocilia are gated by tension. A key regulatory process of MET is *slow adaptation*, thought to enhance the auditory system’s dynamic range. Traditionally, this process has been attributed to myosin motor activity. Here, we challenge this prevailing model and provide evidence for an alternative mechanism in which *phosphatidylinositol 4,5-bisphosphate* (PIP_2_) modulates slow adaptation via interactions with the MET complex protein TMIE. Remarkably, adaptation was rescued by exogenous PIP_2_ even when myosin motors were inhibited, highlighting PIP_2_’s central role. Disruption of TMIE, a PIP_2_-binding protein, also impaired adaptation, and we implicate a PIP_2_ binding site between the channel candidate TMC1 and TMIE to mediate slow adaptation. These findings support a revised model in which PIP_2_–TMIE/TMC1 interactions mediate slow adaptation in hair cells.

## Introduction

The human auditory system is remarkably sensitive, capable of detecting an enormous range of sound intensities and frequencies with extraordinary precision. This ability depends on a series of complex processes that occur within the inner ear, where sound waves are ultimately transformed into electrical signals that the brain can interpret. A key step in this process is known as mechano-electric transduction (MET), the conversion of mechanical energy into electrical activity, which takes place in specialized sensory cells called hair cells^1^. At the top of each hair cell is a bundle of microscopic projections called stereocilia, which bend in response to sound. These movements generate tension in fine extracellular filaments called tip links, which connect adjacent stereocilia and directly control the opening of MET channels. One crucial feature of MET is the system’s ability to adapt, which can act as a protective mechanism that prevents overstimulation and allows the ear to remain sensitive across a wide range of sound levels.

Adaptation appears as a gradual decline in the open probability of MET channels during sustained stimulation. This shift in the channel’s operating range helps the system “reset,” preserving sensitivity to new incoming sounds. Researchers generally describe two types of adaptation^2,3^: a fast component that occurs within milliseconds, and a slow component that unfolds over tens to hundreds of milliseconds. For decades, slow adaptation was thought to rely on myosin motor proteins, particularly myosin Ic (MYO1C)^4,5^, which were believed to climb and slip along the actin filaments in stereocilia while attached to the upper insertion point of the tip link, therefore adjusting tip-link tension. This so-called motor model of adaptation has been the prevailing explanation and is based on studies in non-mammalian systems like the bullfrog sacculus^6–9^ and the mammalian vestibular system^4^. However, our recent work in mammalian hair cells challenges this longstanding model^10^. We previously indicated that the mechanism underlying slow adaptation is not located at the upper insertion point of the tip link, as the motor model suggests, but at the lower end, pointing to a fundamentally different mechanism.

In this study, we explore a different molecular player in slow adaptation: phosphatidylinositol 4,5-bisphosphate (PIP_2_), a lipid molecule known to regulate ion channels. Prior work has implicated PIP_2_ in hair cell function in both bullfrog and mammalian systems, with emerging evidence suggesting a role in adaptation^11,12^. Here, we provide evidence that PIP_2_ is central to slow adaptation in mammalian cochlear and vestibular hair cells. We show that even when myosin motors are inhibited, slow adaptation can be rescued by exogenous PIP_2_, suggesting that the primary role of myosin motors may be in localizing or delivering PIP_2_, rather than directly driving adaptation. Moreover, we examine TMIE, a core MET complex protein, which binds PIP_2_ and is essential for channel function. Using a mouse model (*Tmie^R82C/R82C^*) that disrupts PIP_2_ binding, we demonstrate that disruption of PIP_2_–TMIE interaction impairs slow adaptation, highlighting its critical role in this process. Finally, we identify that TMC1 can also bind PIP_2_ and support a potential PIP_2_ binding pocket between TMIE and TMC1.

Taken together, these findings represent a major step toward redefining the molecular mechanism of slow adaptation in mammalian hair cells, moving beyond the motor model and toward a lipid–protein interaction model with broader implications for understanding mechanosensation.

## Results

### MYO7A-C does not regulate slow adaptation

Myosin VIIA (MYO7A) is an unconventional myosin motor essential for hearing^13,14^, and mutations cause syndromic and non-syndromic recessive hearing loss in humans^15–17^. MYO7A localizes to the upper tip-link density, and this localization places MYO7A in the effective position to serve as the adaptation motor in the previously hypothesized motor model of slow adaptation^7,9,18^. We previously demonstrated that MYO7A is expressed in multiple isoforms: canonical (MYO7A-C), short (MYO7A-S), and N-isoform (MYO7A-N)^19–21^. Specific deletion of the canonical isoform (*Myo7a-ΔC* mouse) severely reduces the levels of *Myo7a* in the inner hair cells (IHCs). MET recordings in *Myo7a-ΔC* IHCs supported a role of MYO7A in generating tip-link tension, but the impact on slow adaptation was not investigated. Here, using the *Myo7a-ΔC* mice, we tested whether slow adaptation is affected. These experiments were performed in IHCs, as outer hair cells (OHCs) from *Myo7a-ΔC* mice showed only limited depletion of MYO7A and no measurable alterations in MET properties. Although IHCs naturally exhibit less slow adaptation than the OHCs (Figure 1B, WT)^10^, we sought to maximize slow adaptation by using an intracellular solution with 0.1 mM BAPTA based on our previous data, which was still insufficient to achieve appreciable slow adaptation magnitude^10^. To increase the magnitude of slow adaptation, we utilized long depolarization modulation (LDM), where we depolarize the cell to +80 mV for ∼10 seconds. This protocol transitions the cell into a state that exhibits greater slow adaptation^22^, which we observe in MET currents after repolarization (Figure 1A, B, WT LDM). We compare WT and *Myo7a-ΔC* IHCs and find that the magnitude of slow adaptation was similar (Figure 1; WT LDM: 29 ± 17%, n = 10; *Myo7a-ΔC* LDM: 26 ± 15%, n = 8; *p* = 0.69, Student’s t-test), indicating MYO7A is not critical for slow adaptation. We did find that resting open probability (resting P_o_) was reduced in *Myo7a-ΔC* IHCs prior to LDM, consistent with previous results and supporting a reduced tip-link tension (Figure 1B)^19^; however, current-displacement (activation) curve differences were not significantly different prior to LDM (Supplemental Figure S1). These results further demonstrate an uncoupling of tip-link tension generation and slow adaptation that we have observed with multiple manipulations^10,22^. This uncoupling and lack of a role of MYO7A in slow adaptation further supports inconsistencies with the motor model of slow adaptation, requiring a new model for slow adaptation.

**Figure 1.**
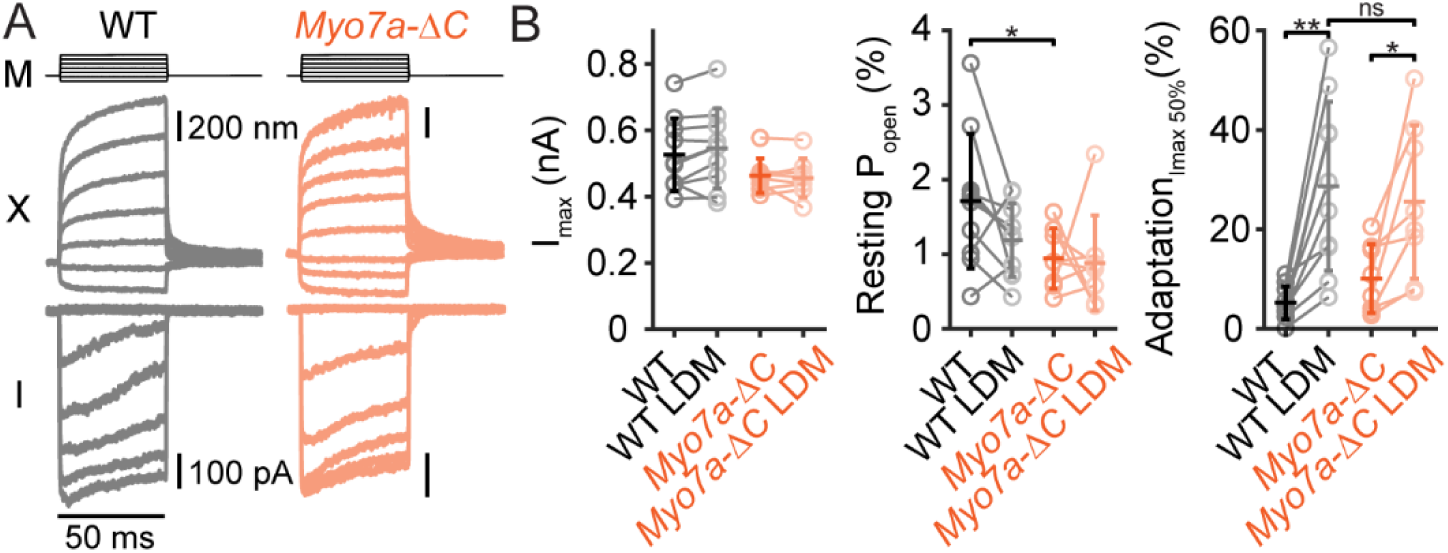
MYO7A does not regulate slow adaptation in cochlea hair cells. **(A)** Example of MET currents (I) and hair bundle displacements (X) from WT (black) and *Myo7a-ΔC* (orange) IHCs elicited by FJ force steps after LDM. **(B)** Summary plots of the peak MET current (I_max_), resting open probability (resting P_o_), and adaptation magnitude measured at 50 % of I_max_. Each plot has WT hair cells before LDM (WT) and after LDM (WT LDM) as well as *Myo7a-ΔC* hair cells before LDM (*Myo7a-ΔC*) and after LDM (*Myo7a-ΔC* LDM). Error bars indicate the mean ± SD. \**p* < 0.05, \*\**p* < 0.01, ns = not significant. Number of cells (animals): Wild type 10 (9), *MYO7A-ΔC* 8 (7).

### PIP2 regulates slow adaptation in cochlear and vestibular hair cells

PIP_2_ is a phospholipid able to regulate multiple cellular processes, including many ion channels^23–25^, and has been shown to regulate the MET channel^11,12,26^. In bullfrog saccular hair cells, PIP_2_ depletion causes a reduction in the rate of fast and slow adaptation as well as reduced MET current^11^. In rat IHCs, PIP_2_ modulates fast adaptation as well as MET channel conductance and open probability^12^, however, slow adaptation in mammalian hair cells was not explicitly tested. Notably, the localization pattern of PIP_2_ suggests a potential role in MET regulation. Therefore, we examined the distribution of PIP_2_ in hair cell stereocilia. PIP_2_ is localized to the tips of both OHC and IHC stereocilia with stronger PIP_2_ antibody labeling at the tips of tallest stereocilia, but labeling is also observed at the tips of shorter stereocilia, albeit at a lower intensity (Supplemental Figure S2A, B)^12^. The labelling of PIP_2_ appears membrane bound, showing a cap-like pattern on the stereocilia (Supplemental Figure S2C). Additionally, staining of PIP_2_ reduces following treatment with phenylarsine oxide (PAO), a PI-4 kinase inhibitor that blocks PIP_2_ synthesis, further supporting the validity of the immunolabelling and the effect of PAO to deplete PIP_2_ (Supplemental Figure S2D).

Having established that PAO effectively depletes PIP_2_ in hair bundles (Supplemental Figure S2D)^12^, we next tested whether PIP_2_ is required for slow adaptation in outer hair cells and utricular hair cells, since both these cell types exhibit robust slow adaptation (Figure 2A, C). We delivered 100 µM PAO extracellularly and measured the change in slow adaptation magnitude in OHCs (Figure 2A, B) and vestibular hair cells (VHCs; Figure 2C, D). After 5 minutes of PAO treatment, the magnitude of slow adaptation in OHCs significantly decreased (Figure 2A, B; Control: 53 ± 18%; PAO: 26 ± 11%; n = 6; *p* = 0.0047, paired Student’s t-test), but not in the DMSO controls (Control: 45 ± 21%; DMSO: 45 ± 19%; n = 8; *p* = 0.93, paired Student’s t-test). We also observed a small decrease in the peak MET current (I_max_) with PAO treatment (Control: 778 ± 57 pA; PAO: 664 ± 119 pA; n = 6; *p* = 0.027, paired Student’s t-test), which is consistent with previous results (Figure 2B)^12^. There were no significant changes observed in the resting P_o_ or activation curve properties (Figure 2B, Supplemental Figure S3). In vestibular hair cells, we observed a decrease in the I_max_, but this was observed in both PAO and DMSO treatment, suggesting this could be due to repeated stimulations causing partial damage to the vestibular hair bundle. These results show that PIP_2_ is critical for slow adaptation in cochlear and vestibular hair cells.

**Figure 2.**
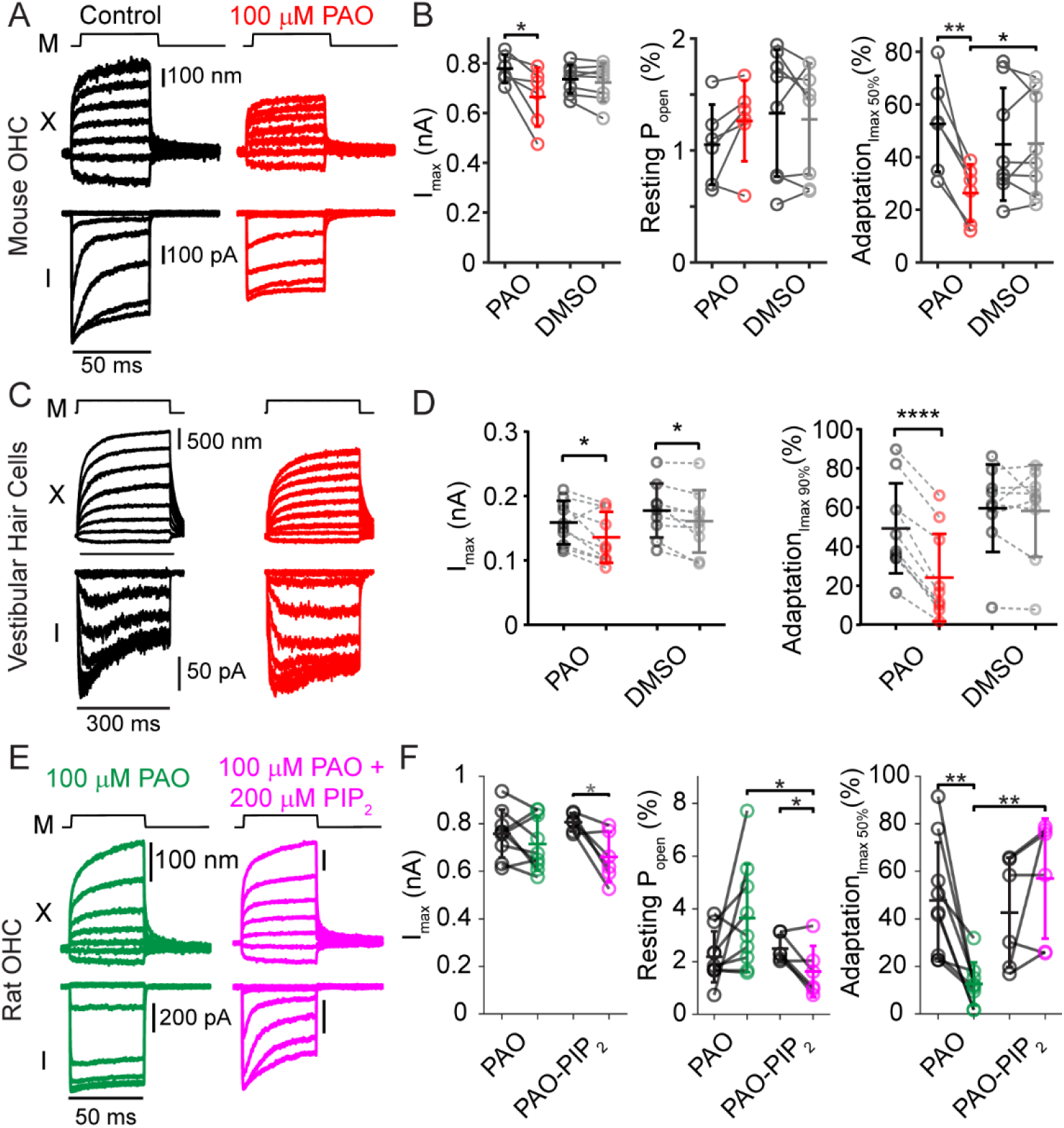
Changes in adaptation in cochlear and vestibular hair cells upon inhibition of PIP_2_ synthesis using PAO. **(A**) Example of hair bundle displacement (X) and MET current (I) recorded from the same mouse OHC before (black) and after 5 minutes of perfusion of 100 µM PAO (red) elicited by FJ force steps (M). **(B)** Summary plots of I_max_, resting P_o_, and adaptation measured at 50% of Imax recorded before and after perfusion of 100µM PAO (PAO). Control cells (DMSO) were recorded before and after 5 minutes of DMSO perfusion. **(C)** Example of hair bundle displacement (X) and MET current (I) recorded from the same type II VHC before (black) and after 5 minutes of perfusion of 100 µM PAO (red) elicited by FJ force steps (M). **(D)** Summary plots of I_max_ and adaptation measured at 90 % of I_max_ recorded before and after perfusion of 100µM of PAO (PAO). Control cells (DMSO) were recorded before and after 5 minutes of DMSO. **(E)** Example of hair bundle displacement (X) and MET current (I) recorded in rat OHCs after perfusion of PAO with normal intracellular solution (green) or intracellular solution supplemented with 200 µM PIP_2_ (magenta). **(F)** Summary plots for rat OHCs with normal intracellular solution or supplemented with 200 µM PIP_2_ before and after PAO treatment are shown like panel B. Pre-and post-PAO/DMSO treatment for individual cells are connected. Error bars indicate the mean ± SD. \**p* < 0.05, \*\**p* < 0.01, \*\*\*\**p* < 0.0001. Number of cells (animals): mouse OHCs with PAO 6 (6), mouse OHCs with DMSO 9 (9), VHCs with PAO 10 (10), VHCs with DMSO 9 (9), rat OHCs with PAO 9 (9), rat OHCs with PAO and PIP_2_ 6 (6).

To further confirm the importance of PIP_2_ and ensure that PAO is indeed acting on PIP_2_ levels in the hair cell, we performed an experiment where we included PIP_2_ in the intracellular solution. Since PAO reduces the synthesis of PIP_2_, exogenous PIP_2_ supplementation could presumably reduce the depletion of PIP_2_ in the hair cells. For this experiment we decided to also extend the findings to other species and performed the rescue experiment using rat OHCs. We found that in control conditions without intracellular PIP_2_, PAO reduced the magnitude of slow adaptation in rat OHCs similar to our results in mouse OHCs (Figure 2E-F, green; *p* = 0.003); however, when PIP_2_ was included in the intracellular solution, slow adaptation was not significantly reduced upon PAO application (Figure 2E-F, magenta; *p* = 0.19). Peak MET current and resting P_O_ changes were not consistent with changes observed in mouse OHCs, therefore, less emphasis is placed on any differences observed. Additionally, the magnitude of slow adaptation after PAO treatment was significantly higher when PIP_2_ was in the intracellular solution (Figure 2F; *p* = 0.006). These results show that inclusion of intracellular PIP_2_ can rescue the effect of PAO, further supporting that depletion of PIP_2_ is the driver of the reduced slow adaptation during PAO treatment.

Since a hallmark of slow adaptation is a shift in the activation curve of the channel, we tested the effects of PAO on shifts of the activation curve by using a multi-pulse protocol (Figure 3A)^27^. In this protocol, an activation curve is generated before an adapting force step is applied. Then 10 ms and 50 ms after the application of an adapting force step, activation curves are generated to determine the adaptive shifts in the activation curve (Figure 3B). With PAO treatment, the shifts in the activation curve significantly decreased between the 1^st^ and 3^rd^ activation curves and the 2^nd^ and the 3^rd^ activation curves (Figure 3C; Δ ½ Act_3-1_, *p* = 0.038; Δ ½ Act_3-2_, *p* = 0.011), indicating that adaptation was reduced with PAO treatment. Importantly, this reduction is not apparent in the control case (0.1% DMSO perfusion; Δ ½ Act_3-1_, *p* = 0.94; Δ ½ Act_3-2_, *p* = 0.79) or when PIP_2_ supplemented in the intracellular solution with PAO treatment (magenta; Δ ½ Act_3-1_, *p* = 0.32; Δ ½ Act_3-2_, *p* = 0.20), suggesting PIP_2_ depletion is the cause of the reduced adaptation observed. For the post-treatment condition, the shifts in the activation curve were less in the PAO treatment as compared to the DMSO treatment (Δ ½ Act_3-1_, *p* = 0.034; Δ ½ Act_3-2_, *p* = 0.016). The resting properties of the MET channel (i.e. 1^st^ activation curve in the multi-pulse protocol) were not significantly different from each other before and after application of PAO (Supplemental Figure S4), which is similar to the results found when using a force step paradigm (Supplemental Figure S3). These data indicate that PIP_2_ depletion with PAO does not significantly alter the resting state properties of the activation curve, but significantly affects the adaptation mechanism itself therefore affecting activation curve properties only after a stimulus is applied. This behavior is consistent with PIP_2_’s role occurring after stimulus application to mediate the adaptation process.

**Figure 3.**
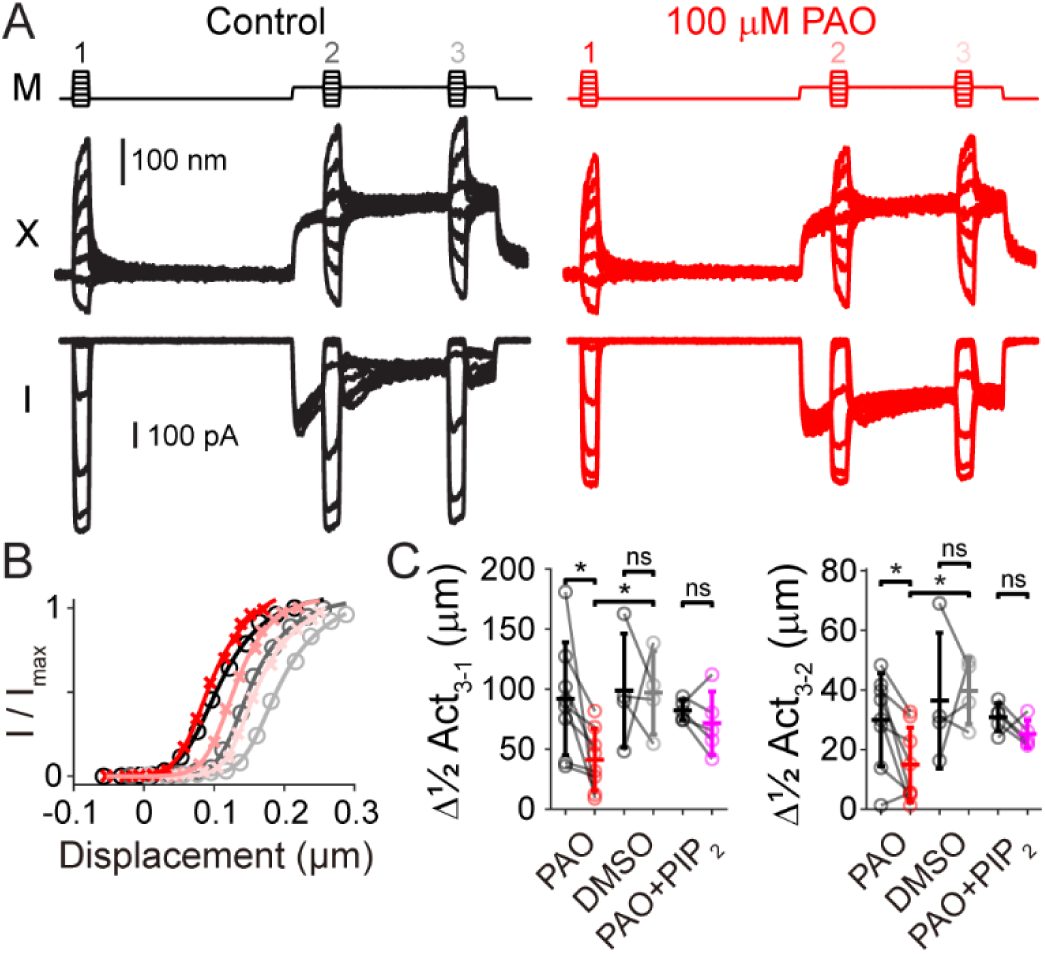
Multi-pulse protocols indicate reduced shifts in the presence of PAO. **(A)** Example of hair bundle displacement (X) and MET current (I) elicited by fluid-jet force waveforms (M) from a P8 rat OHC using intracellular solution containing 0.1 mM BAPTA before (black) and after perfusion with 100 µM PAO (red). **(B)** Resulting activation curves for each condition with each activation curve progressively shifting over time. Color coding matches number colors above stimulus waveform (M) in panel A. **(C)** Summary of the shifts in P7-P8 rat OHCs between the ½ activation for pulse 3 and pulse 1 (left) ad pulse 3 and pulse 2 (right) for before (black) and after perfusing 100 µM PAO (red, n=8), 0.1% DMSO (gray, n=4), and 100 µM PAO with 200 µM PIP_2_ supplemented in the intracellular solution (magenta, n=5). Error bars indicate the mean ± SD. * *p* < 0.05.

### Myosin motors have a secondary role in the regulation of slow adaptation

Myosin motors are important for slow adaptation in hair cells^28,29^. Specifically, MYO1C regulates slow adaptation in vestibular hair cells^4,10^, and we find that PIP_2_ is also concentrated in the stereocilia tips in vestibular hair cells (Supplemental Figure S2E). In the previously proposed motor model of adaptation, myosin motors were thought to play a primary role in slow adaptation by releasing tension at the upper tip-link insertion^7,9^. However, the aforementioned evidence against the motor model challenges this interpretation and raises the question of how myosin motors actually participate in slow adaptation^10^. If PIP_2_ directly regulates slow adaptation, myosin motors may have an indirect role in slow adaptation, potentially involving their well-established function in cargo transport. MYO1C is able to bind PIP_2_^30,31^, and we hypothesize that MYO1C may transport and/or concentrate PIP_2_ close to the MET channel to regulate slow adaptation. We used the *Myo1c^Y61G/Y61G^* mouse model (Figure 4A) to test this hypothesis and blocked MYO1C Y61G activity by adding intracellular N^6^(2-methyl butyl) Adenosine Diphosphate (NMB-ADP), an allele specific MYOC1C inhibitor (Figure 4B). In agreement with prior studies, inhibition of MYO1C Y61G with NMB-ADP reduced the magnitude of slow adaptation (control: 36 ± 23%, n = 25; NMB-ADP: 19 ± 9%, n = 13; *p* = 0.002, Student’s t-test)^4,5,10^. We next tested whether exogenous PIP_2_ could rescue the magnitude of slow adaptation (Figure 4C). Remarkably, inclusion of PIP₂ in the intracellular solutions with NMB-ADP maintained slow adaptation at control levels (36 ± 22%, n = 11; *p* = 0.95, Student’s t-test vs *Myo1c^Y61G/Y61G^*; Figure 4C, D). Slow adaptation time constants were 90 ± 38 ms, n = 17 in control and 94 ± 28 ms, n = 7 in PIP_2_ rescued cells (*p* = 0.79), suggesting similar adaptation processes in both control and rescued conditions. NMB-ADP inhibition in *Myo1c^Y61G/Y61G^*reduces I_max_^4,5,10^, and we found this decrease was also rescued with exogenous PIP_2_ in the intracellular solution (Figure 4D), further supporting rescued MET function. These results demonstrate that exogenous PIP_2_ is sufficient to preserve slow adaptation when myosin motor activity is inhibited, indicating an indirect role of MYO1C in the mechanism of slow adaptation. This indirect role of myosin motors is further supported by Figure 2 that showed a decrease in slow adaptation when PAO inhibited PIP_2_ synthesis presumably without myosin motor inhibition, indicating myosin motor activity itself is not sufficient for slow adaptation. These striking data together indicate that PIP_2_ is likely downstream of myosin motor function in the slow adaptation mechanism as PIP_2_ can functionally compensate for myosin motor inhibition.

**Figure 4.**
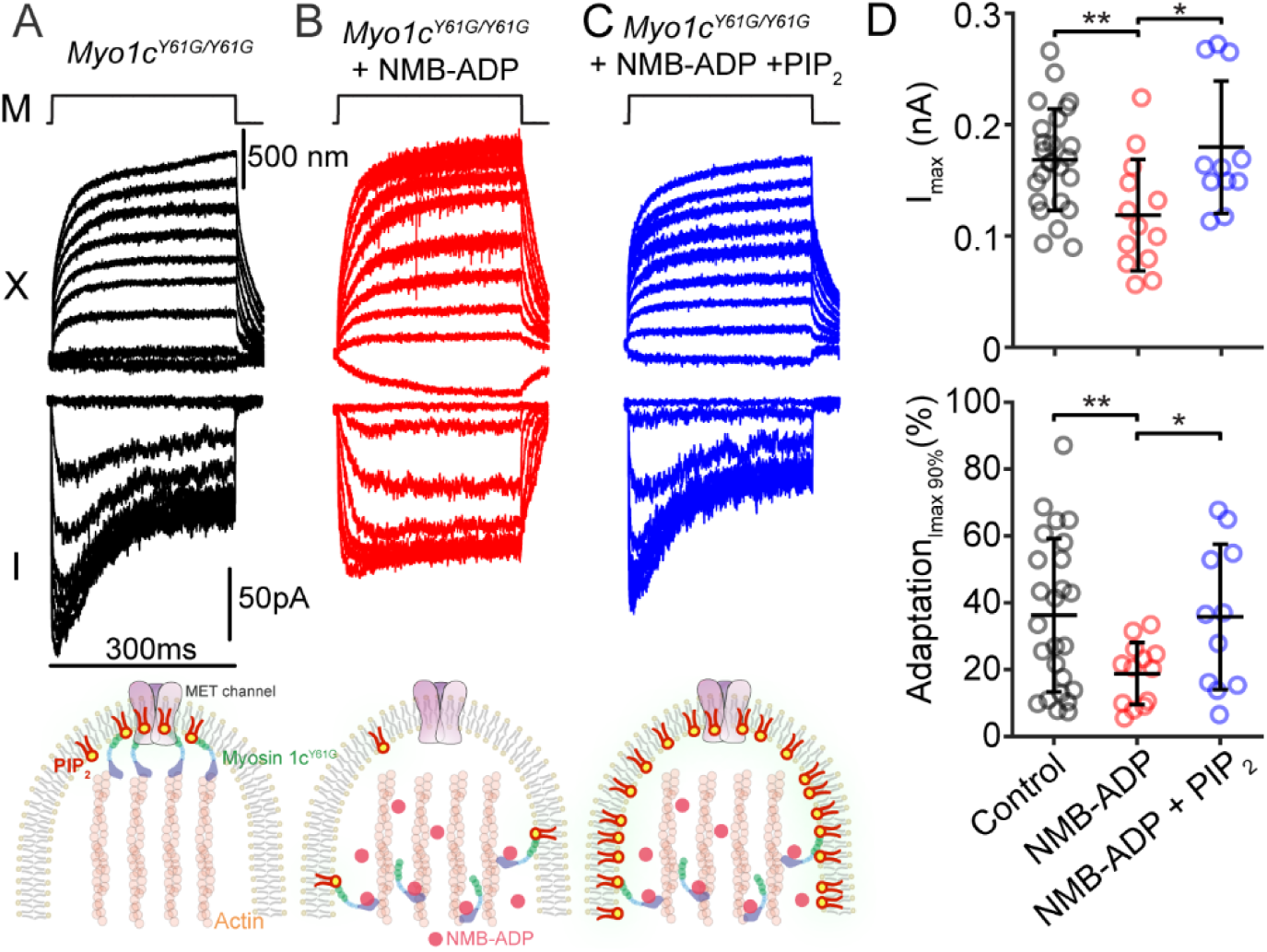
PIP_2_ restores slow adaptation in VHCs after inhibiting MYO1C activity. **(A – C)** Example of hair bundle displacement (X) and MET current (I) recorded in **(A)** *Myo1c^Y61G/Y61G^* type II VHCs in 0.1 mM BAPTA, **(B)** 250 µM NMB-ADP, and **(C)** 250 µM NMB-ADP + 25 µM PIP_2_ elicited by fluid-jet force steps (M). Corresponding schematics of the hypothesized underlying mechanism (bottom): **(A)** control condition in *Myo1c^Y61G/Y61G^*hair cells, MYO1C is able to transport/concentrate PIP_2_ close to the MET channel **(B)** inhibition of MYO1C resulted in less PIP_2_ located in the proximity of the MET channel, and **(C)** inhibition of MYO1C with exogenous PIP_2_ is able to rescue slow adaptation by increasing the overall concentration of PIP_2_ in the membrane, including near the MET channel. **(D)** Summary plots of peak current (top) and adaptation (bottom) measured at 90% of I_max_. Error bars indicate the mean ± SD. * *p* < 0.05, ** *p* < 0.01. Number of cells (animals): *Myo1c^Y61G/Y61G^* 25 (21), *Myo1c^Y61G/Y61G^* with 250 µM NMB-ADP 13 (10), *Myo1c^Y61G/Y61G^* with 250 µM NMB-ADP and 25 µM exogenous PIP_2_ 11 (9). Some of the cells for control and NMB-ADP were included from^10^.

In contrast to vestibular hair cells, inhibition of MYO1C in the *Myo1c^Y61G/Y61G^* mouse model does not lead to a reduction in slow adaptation magnitude in cochlear hair cells^10^. We therefore employed an alternative method to disrupt myosin motor function and determined whether exogenous PIP_2_ could rescue slow adaptation in cochlear hair cells. We and others previously showed that intracellular sulfate (SO_4_^2-^), which inhibits myosin motor function, reduces the slow adaptation magnitude in OHCs^8,10^ (Figure 5A, B), and we used this paradigm to test exogenous PIP_2_ rescue of slow adaptation. Similar to previous results, we observed a significant reduction of slow adaptation magnitude when SO_4_^2-^ was included in the intracellular solution (Figure 5D, black to red data points; control: 54 ± 23%, n = 7; SO_4_^2-^: 12 ± 8%, n = 12; *p* = 0.002, Student’s t-test). To determine whether PIP_2_ could rescue this effect, we tested multiple intracellular PIP_2_ concentrations. While 100 µM PIP_2_ provided partial rescue (Figure 5D, light blue; 25 ± 24%, n = 12 for 100 µM PIP_2_), a higher concentration (200 µM PIP_2_) significantly increased slow adaptation, restoring it to near control levels (Figure 5D, blue data points; 52 ± 24%, n = 8; *p* = 0.002, Student’s t-test) (Figure 5C, D). Slow adaptation time constants were 10 ± 2 ms (n = 7) in control and 12 ± 8 ms (n = 8) in 200 µM PIP_2_ rescued cells (*p* = 0.51, Student’s t-test), supporting that the adaptation process was similar. We also observed a slight increase in resting P_o_ with SO_4_^2-^inhibition that was also rescued with PIP_2_ (Figure 5D), however, this difference was not reflected in the overall activation curve properties (Supplementary Figure S4). While we cannot fully rule out the possibility that exogenous PIP_2_ rescues slow adaptation through an off-target effect, such as interfering with the inhibition of the myosin motor blocker, this seems unlikely because the rescue was effective in two different hair cell types using distinct myosin motor inhibition strategies. Together, these results reinforce the conclusion that PIP_2_ plays a central and direct role in regulating slow adaptation, with myosin motors contributing indirectly and upstream of the role of PIP_2_.

**Figure 5.**
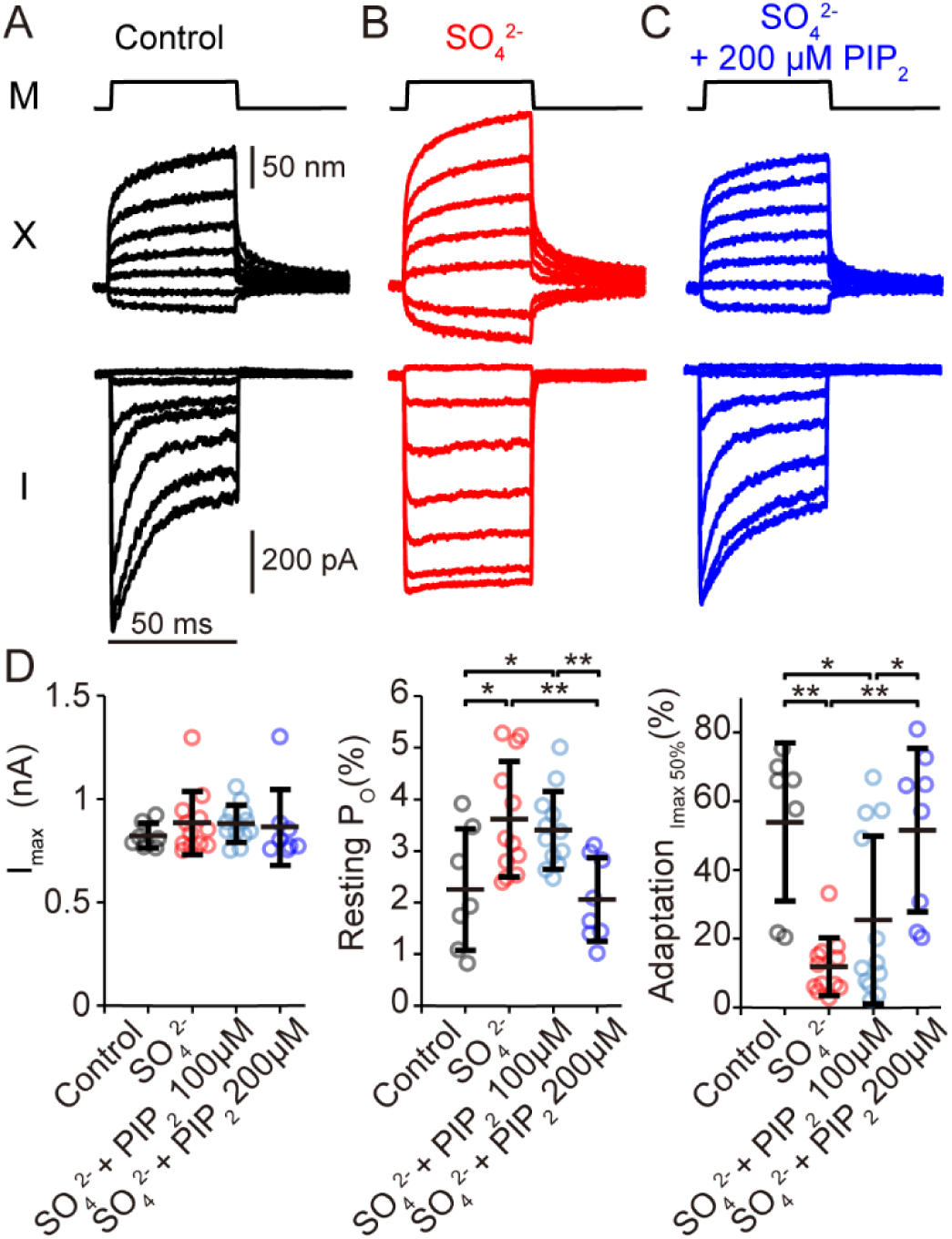
PIP_2_ restores slow adaptation in OHCs after inhibiting myosin ATPase activity. **(A - C)** Example of hair bundle displacement (X) and MET current (I) elicited by fluid-jet force steps (M) from P6-7 rat OHCs using intracellular solution containing **(A)** 0.1 mM BAPTA, **(B)** 0.1 mM BAPTA with 60 mM SO_4_^2-^ (red), and **(C)** 0.1 mM BAPTA with 60 mM SO_4_^2-^ and 200 µM PIP_2_ (blue) elicited by fluid-jet force steps. **(D)** Summary plots for different intracellular solutions of the I_max_, resting P_O_, and adaptation magnitude. Error bars indicate the mean ± SD. \**p* < 0.05, \*\**p* < 0.01. Number of cells (animals): Control 7 (7), SO_4_^2-^ 12 (12), SO_4_^2-^ + PIP_2_ (100 µM) 12 (12), SO_4_^2-^ + PIP_2_ (200 µM) 8(8).

### TMIE is involved in slow adaptation

TMIE is a protein that is imperative for hair cell MET^3,26,32^. PIP_2_ also binds TMIE based on PIP strip binding assays, and mutation of key arginine residues inhibits PIP strip binding^26^. In hair cells, depletion of PIP_2_ was shown to reduce the MET current faster in *Tmie^R82C/R82C^* vs *Tmie^R82C/+^*, suggesting that the binding of PIP_2_ is reduced with the R82C mutation^26^.

Given our hypothesis that slow adaptation requires PIP_2_, we tested whether TMIE might influence slow adaptation by mediating a PIP_2_ interaction with the MET channel complex. We used the *Tmie^R82C/R82C^* mice and control heterozygotes (*Tmie^R82C/+^*) to measure differences in slow adaptation. Experiments were conducted at P4-P5, when MET currents are still present in *Tmie^R82C/R82C^*, though with reduced amplitude (Figure 6A, C)^26^. At this age, adaptation was less apparent in control *Tmie^R82C/+^* mice, so we augmented slow adaptation again by using LDM^22,33,34^. We observed a shallowing of the activation curve in *Tmie^R82C/R82C^* hair cells that corresponds with the observed increased resting P_o_ of the channel (Figure 6C and Supplementary Figure S5). This phenomenon was not previously reported and may be due to using a fluid-jet stimulation as compared to a stiff probe stimulation^26^. In control *Tmie^R82C/+^*mice, we observed robust slow adaptation (32 ± 16%, n = 18), but in *Tmie^R82C/R82C^* mice, slow adaptation was significantly reduced (11 ± 10%, n = 20; *p* = 0.000021, Student’s t-test) (Figure 6A, C). This data indicates a potential role of TMIE in slow adaptation.

**Figure 6.**
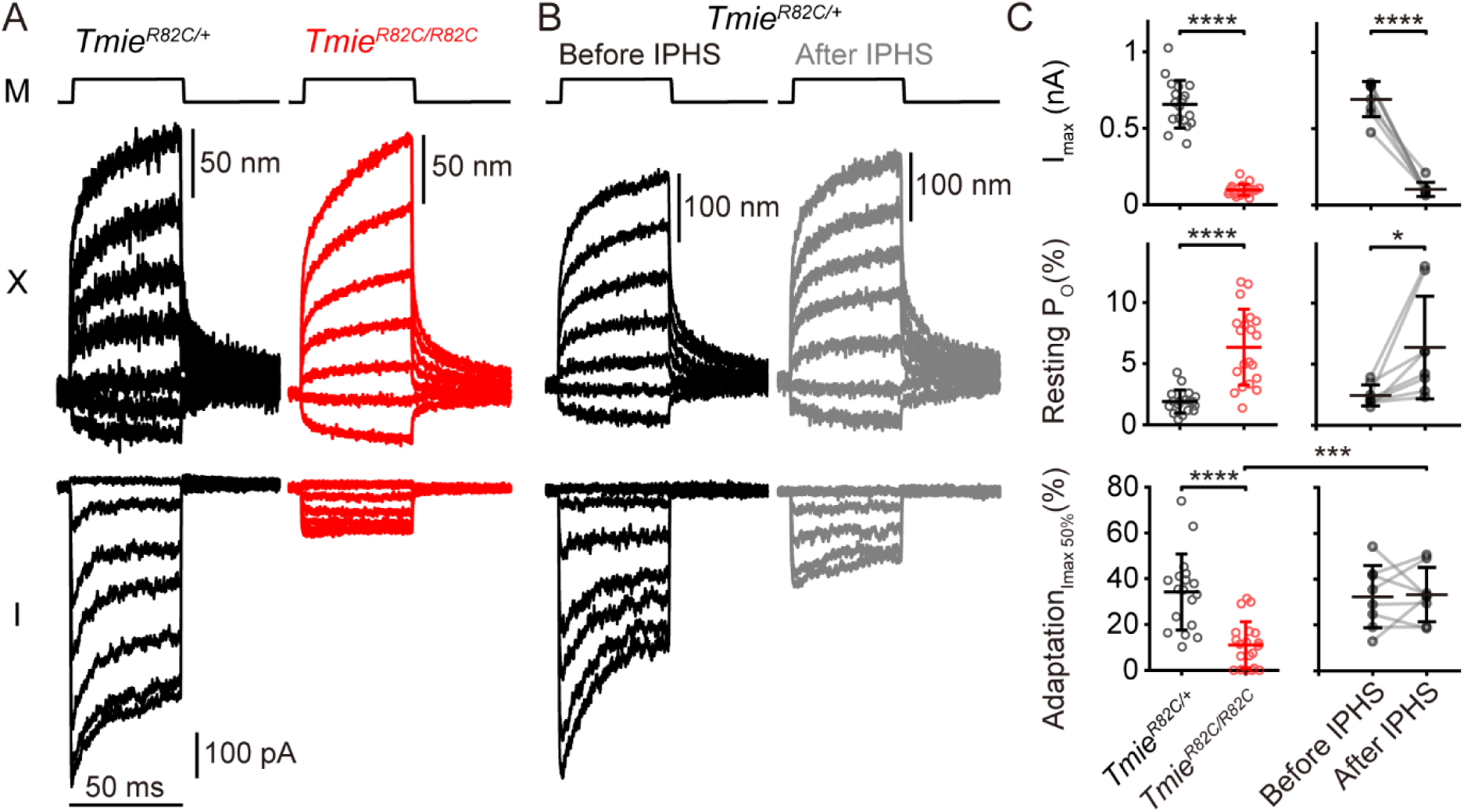
TMIE (R82C) mutation results in reduced slow adaptation that is not a due to the reduced MET current. **(A)** Displacement (X) and current (I) measurements elicited by fluid-jet force steps (M) on cochlear outer hair cells form P4-P5 *Tmie^R82C/+^* and *Tmie^R82C/R82C^* mice. Using an intracellular solution containing 0.1 mM BAPTA, the *Tmie^R82C/+^* (black trace) and *Tmie^R82C/R82C^* (red trace) were analyzed by applying a step-like force stimulus at -80 mV following a 10 sec depolarization at + 80 mV. **(B)** To determine whether the lowered MET current in *Tmie^R82C/R82C^*affects slow adaptation, IPHS was performed using 100 mM EDTA to reduce the MET current in *Tmie^R82C/+^* hair cells. In *Tmie^R82C/+^*, measurements of the displacement (top) and I_max_ (top) using a step-like force stimulus at -80 mV after 10 sec depolarization at +80 mV were shown as black traces before IPHS and gray traces after IPHS. **(C)** Summary plots of data show the peak current, percentage of resting P_o_, and slow adaptation magnitude. Slow adaptation was measured at the 50 % point of the total peak current. Error bars indicate the mean ± SD. * *p* < 0.05, *** *p* < 0.001, **** *p* < 0.0001. Number of cells (animals): *Tmie^R82C/+^* 18 (18), *Tmie^R82C/R82C^* 20 (20), IPHS *Tmie^R82C/+^* 8 (8).

One potential confounding factor is the reduction in I_max_ in *Tmie^R82C/R82C^* hair cells (Figure 6A, B)^26^, which could impact Ca^2+^ influx, a key driver of slow adaptation. To test whether reduced current alone could account for the adaptation deficit, we mimicked the decreased current of *Tmie^R82C/R82C^* mice in *Tmie^R82C/+^* controls by breaking a subset of tip links with EDTA iontophoresis (IPHS), which lowers MET current amplitude without altering TMIE protein^35–37^. After IPHS, *Tmie^R82C/+^* cells displayed I_max_ comparable to those of *Tmie^R82C/R82C^* (*Tmie^R82C/R82C^* : 96 ± 37 pA, n = 20; *Tmie^R82C/+^* after IPHS: 102 ± 46 pA, n = 8; *p* = 0.73, Student’s t-test)^22,37^. Despite the reduced current amplitudes, the slow adaptation magnitude did not change significantly after IPHS (Figure 6C; *p* = 0.89, paired Student’s t-test), and the slow adaptation magnitude was also significantly higher than *Tmie^R82C/R82C^* hair cells (*p* = 0.0006, Student’s t-test). These data suggest that the reduced slow adaptation magnitude in *Tmie^R82C/R82C^* OHCs was not due to the reduced I_max_ and support a role of TMIE in the slow adaptation process.

### PIP2 can interact with TMC1 and TMIE in the MET complex

The MET complex is hypothesized to contain at least TMC1, TMIE, and CIB2 ^3,32,38–41^. The cryo-EM structure of native *C. elegans* TMC-1 was solved in complex with hair cell MET component homologs of TMIE and CIB2^42^. Using this structure our group previously modeled the mammalian MET complex^43^. The location of TMIE R82 sits at the interface between TMC1 and TMIE, raising the possibility that PIP_2_ may interact with the MET complex in the region between TMC1 and TMIE. Previous work already demonstrated that TMIE could directly interact with PIP ^26^, so we performed similar experiments to determine if TMC1 could also directly interact with PIP_2_. Using a lipid strip assay, we found that purified mouse TMC1 was able to bind to phosphoinositide binding spots, with PIP_2_ (PI(4, 5)P_2_) binding as well, albeit at lower levels as compared to some other phosphoinositide lipids (Fig 7A, B). PIP_2_ specific binding with a PIP_2_ grip demonstrated PI(4,5)P_2_ specificity while a blank spot lacking lipids served as a control for potential non-specific binding. These data support that TMC1 can also bind PIP_2_, supporting a hypothesis that both TMC1 and TMIE directly interact with PIP_2_.

**Figure 7.**
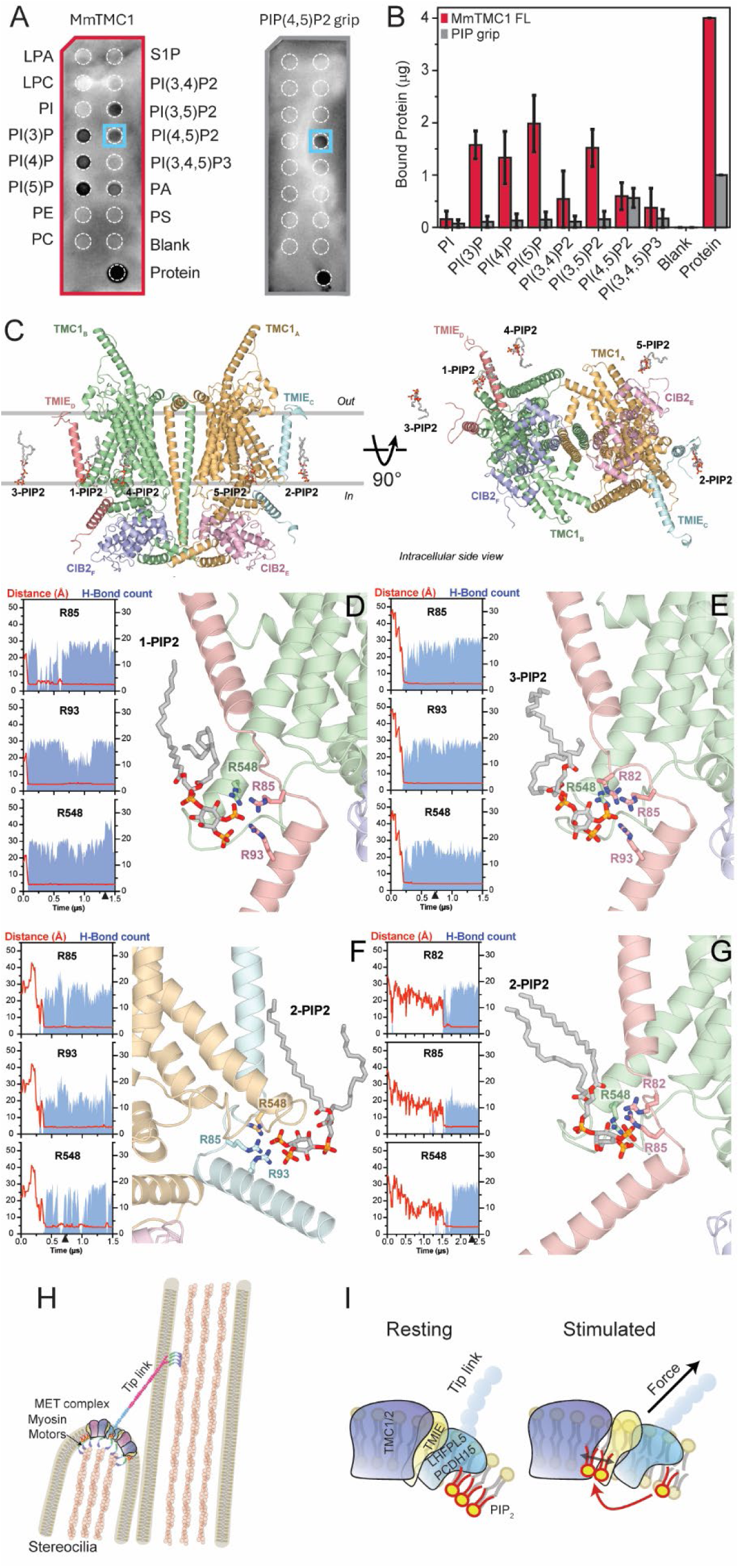
An MET complex binding pocket for PIP_2_ with proposed new schematic model of slow adaptation. **(A)** Purified full-length MmTMC1 directly interacts with phosphoinositide lipids on PIP strip blots (left). PI(4,5)P2 binding is indicated with a blue square. Positive control PIP_2_ grip protein only interacts with PIP_2_ (right blot). **(B)** Quantification of the data shown in B (mean ± SEM from 3 (MmTMC1) and 2 (PIP_2_ grip) independent experiments). **(C)** Schematic of the MET complex used in molecular dynamics simulations at time 0 with side view (top) and rotated viewed from the intracellular side (bottom). **(D-G)** Simulation results from the three replicas that had an interaction of PIP_2_ with residues R82 or R85 of TMIE. For each simulation replica, overlay plots (left) show the minimum instantaneous distance (d(t)) between each arginine (Cζ - carbon of the guanidil group) and the nearest PIP_2_ phosphate group (d(t) = min[d_P4_(t), d_P5_(t)]), averaged over 10 ns bins (red trace, left axis), alongside the corresponding H-bond count per bin (blue shaded area, right axis). All MD simulations showed a stabilization of the PIP_2_ in the binding site. One replica had PIP_2_ molecules interacting with both TMC1 subunits (D and F). Structures shown correspond to the timepoint noted by the arrowhead. Stick representation of residues on TMC1 and TMIE are shown that have significant interactions in the form of hydrogen bonds with PIP_2_. **(H)** Schematic of two adjacent stereocilia illustrates some core components of the MET complex. This model highlights the key role of PIP_2_ (red lipids) in slow adaptation, suggesting that myosin motors are important only for transport/localization of PIP_2_. **(I)** An enlarged view showing half of the MET complex illustrating that tip-link force may uncover a binding site for PIP_2_ between TMC1/2 and TMIE to mediate slow adaptation when the channel complex is stimulated.

To investigate where PIP_2_ may bind to the MET complex and assess the stability of the interaction, we performed six independent molecular dynamics (MD) simulations of the MET complex embedded in a POPC membrane containing PIP_2_ at 2% of the inner leaflet phospholipids (Figure 7C). Since the negatively charged headgroup of PIP_2_ typically binds to positively charged amino acids, we found residue TMC1 R548 that was in the vicinity of TMIE R82 as well as TMIE R85 and R93, both of which were also identified previously as important residues for PIP_2_ binding^26^. We equilibrated the system and looked at whether PIP_2_ would bind in this TMIE-TMC1 region of the MET complex. In all replicas, the arginine residues R85 TMIE and R548 TMC1 (as well as R82 TMIE and R93 TMIE where monitored) initially resided at distances of 30–50 Å from the PIP_2_ phosphate groups and transitioned to stable contact (Cζ–P distances of approximately 4–6 Å) in three of the six replicas (Figure 7D-G; Supplementary Video 1; Supplementary Figure S7). Once established, these contacts were maintained for the remainder of each simulation (at least 0.5 µs), as indicated by the concurrent and sustained elevation of the H-bond count (Fig. 7D–G, blue shaded areas). The number of H-bonds per 10 ns bin ranged from ∼10 to 30 contacts per residue across replicas, reflecting a robust electrostatic engagement between the guanidinium groups of the arginine side chains and the polyanionic phosphate head groups of PIP_2_. Collectively, these results demonstrate that the PIP_2_-arginine interaction network at the TMC1/TMIE interface is a consistent and reproducible feature of the MET complex, supporting its functional relevance for channel regulation.

## Discussion

With previous models of slow adaptation no longer fitting with current experimental data^10^, we propose and support a new model of slow adaptation where PIP_2_ interacts with TMIE and TMC1 to mediate the slow adaptation process regulating hair cell MET channel activity. Our experimental data continues to draw a distinction between the slow adaptation mechanism and resting tension control in the hair cell MET complex (Figure 1). We also find that the mammalian slow adaptation mechanism requires PIP_2_ (Figure 2), but not the continuous activity of myosin motors (Figure 4, 5). These data indicate that PIP_2_ may directly modulate the MET channel to mediate the slow adaptation process. Others have shown that PIP_2_ and TMIE interact^26^, and our data support that this interaction is likely important for the slow adaptation process, indicating TMIE’s involvement in slow adaptation. Finally, our simulations demonstrate a binding pocket that exists between TMC1 and TMIE that is likely critical for the slow adaptation process (Figure 7). Overall, we support a new model of slow adaptation where PIP_2_ interacts with the MET complex to mediate slow adaptation (Figure 7H, I).

### Slow adaptation mechanism location

Slow adaptation has long been known to be reliant on Ca^2+ 2,6,10^. When MET channels were localized to the lower tip-link region^44^, Ca^2+^ was thought to either diffuse farther away from the channel itself to mediate slow adaptation at motors at the upper tip-link density, or be regulated by bulk Ca^2+^ levels in the stereocilia^33,45^. Based on the mechanical properties of the hair bundle not correlating with slow adaptation, we proposed that the mechanism of slow adaptation originates near the MET channel itself, rather than relying on diffusion to a distant site from the channel^10^. If regulation occurred at the upper tip-link insertion site, then modulating the MET channels located at the lower tip-link insertion would necessitate regulation through the tip link; this would require a mechanical change in the hair bundle associated with slow adaptation in order to reduce force transmission to the MET channel^10^. This is not consistent with our result that reduction of the candidate tip-link motor, MYO7A, did not affect adaptation (Figure 1), further supporting that the slow adaptation mechanism is not at the upper tip-link region.

Further support for our channel modulation model comes from the data presented in Figure 6B, which show that reducing total MET current via tip-link breakage, thereby reducing Ca²⁺ influx which would reduce bulk intracellular free Ca²⁺ concentration, did not significantly diminish the magnitude of slow adaptation (Figure 6C). If Ca²⁺ acted at a distant site from the channel, such a reduction in total Ca²⁺ influx into the hair bundle would be expected to impair adaptation. The absence of such an effect strengthens the idea that Ca²⁺ acts locally at the channel site to mediate adaptation. This conclusion is also consistent with studies in other labs showing slow adaptation-like processes in single-channel ensemble currents, where only the local Ca²⁺ concentration near the open channel would be elevated during stimulation^46,47^. Finally, our findings that TMIE is required for normal slow adaptation further support this model. Given TMIE’s close association with the MET complex, its role in regulating adaptation (Figures 6 and 7) reinforces the conclusion that slow adaptation is tightly coupled to, and likely occurs at, the MET channel complex itself.

### Myosin Motor Role

Myosin motor activity is known to be critical for slow adaptation. Early studies showed that ATPase blockers inhibit slow adaptation^8^, and the localization of MYO1C in the hair bundle positioned it as a strong candidate for the adaptation motor^48^. Definitive support came from experiments using a modified MYO1C Y61G allele, which enables specific inhibition by the NMB-ADP analog. These studies confirmed that MYO1C activity was required for slow adaptation in vestibular hair cells^4,5,10^. While this body of work continues to support a role of MYO1C in slow adaptation, our ability to rescue the reduction of slow adaptation during MYO1C inhibition by supplementing PIP_2_ suggests an indirect role of the motor (Figure 4). Based on this data, we hypothesize that MYO1C is involved in the transport or concentration of PIP_2_ near the MET channel where PIP_2_ interaction with the channel complex drives the slow adaptation process. Thus, when MYO1C activity is reduced, then PIP_2_ is not available to regulate the MET channel complex to mediate slow adaptation.

The data presented here suggest that multiple myosin motor populations serve different roles in hair cell MET. Based on previous models of MET, MYO7A which is localized at the upper tip-link density and is involved in tensioning of the tip link, would have been thought to be involved in slow adaptation as well, since the mechanisms of tensioning and slow adaptation were thought to be the same^9^. However, we found that adaptation is unaffected in MYO7A mutants that exhibit reduced tip-link tension (Figure 1)^19^. The separation of functions allows more points of intervention and control of the MET process. In vestibular hair cells, we speculate that MYO1C transports and/or concentrates PIP_2_ for PIP_2_ to mediate slow adaptation near the lower end of the tip link, but presumably MYO7A would be important for the tension generation at the upper end of the tip link. The separation of functions in this case is by localization, but differences in functions can also exist at different time points in the lifecycle of the cell. For instance, MYO7A function early in development is required for proper hair bundle morphology and later required for proper MET function^14,19^. Recent work suggests that MYO7A may have a third function in adult hair cells^49^. Multiple myosin isoforms may have different functions both at specific locations as well as at specific times.

### Lipid roles in hair cell MET

There is increasing interest in lipid roles in hair cell MET. Early work on PIP_2_ showed the importance of lipids on channel properties, but originally ascribed them to effects on slow adaptation and potentially modifying channel conductance^11^. Later work demonstrated that lipid modulation could affect the gating of the hair cell MET directly via modulation by GsMTx4^34^. Original models assumed direct tethering of the tip-link to the MET channel to mediate activation^9,50^, but newer models incorporate lipids in a potential activation mechanism^51^. MET channels also exhibit lipid scrambling activity^52,53^, furthering the importance of the lipid membrane in MET. Recent work also indicates that putative MET channels composed of TMC1 can be intrinsically activated by lipid stretch and deformation without tip links^54–56^. It is possible that although the channels themselves can be intrinsically activated by the lipid membrane, proteins in the MET complex like LHFPL5 augment the MET channel sensitivity by also creating some direct tethers^57–59^. Nonetheless, if channels can be intrinsically activated by lipid tension, then they should be modulated by the surrounding lipid environment, consistent with the force-from-lipids model of MET^60^. Our data further support this notion, where PIP_2_ has a direct role in the slow adaptation mechanism. Slow adaptation effects are just one way that PIP_2_ regulates the hair cell MET channel. PIP_2_ also affects channel conductance and fast adaptation of hair cell MET and activation of the TMC1-related Ca^2+^-activated chloride channel TMEM16A^11,12,26,61–63^. This raises the likelihood that there may be multiple PIP_2_ binding sites within the MET channel complex that affect channel properties in different ways.

A potential model of how PIP_2_ may mediate slow adaptation is through direct channel modulation. That the activation curves before and after PIP_2_ depletion by PAO remain largely the same (Supplementary Figures S3, S4), suggests that PIP_2_ has little effect on the MET channel prior to any stimulation. This would imply that PIP_2_ action (i.e., PIP_2_ binding to the channel complex) is required for the changes associated with slow adaptation, not for the resting physiological state. TMC1 and TMC2 are structurally similar to TMEM16A^42,64–66^, and TMEM16A has a known regulation by Ca^2+^ as well as PIP_2_^61–63^. The R82C TMIE mutation affecting slow adaptation is located near lipid densities in the *C. elegans* TMC-1 and TMC-2 structures between TMIE and TMC1/TMC2^42,66^ and we have found in molecular dynamics simulations that this location can bind PIP_2_ (Figure 7). The homologous residue to mammalian R82 is *C. elegans* TMIE R49, and this residue makes hydrogen bonds with backbone carbonyl atoms in TMC-1 TM6 in the *C. elegans* structure. We speculate that PIP_2_ may displace this hydrogen bond and change its conformation to create interactions between TMIE, TMC1, and PIP_2_. The binding of PIP_2_ in this pocket (Figure 7D-G) could then lead to what is observed as slow adaptation. With the location of this potential binding pocket being located between TMIE and TMC1, one could envision that force on the MET complex lowers the energy barrier to break the hydrogen bond between TMIE and TMC1, which then reveals the binding site for PIP_2_ to bind the channel and change the energy landscape of the channel, resulting in adaptation (Figure 7H, I).

Not yet addressed is that we know that Ca^2+^ has a role in slow adaptation as well, and its exact mechanism of action requires further investigation. It is possible that interactions of PIP_2_ with the channel are facilitated by calcium or that calcium is a required co-factor for the required conformational change. The role of calcium in this mechanism still requires further elucidation, but the work presented here supports the critical requirement of PIP_2_.

Our description of the roles of PIP_2_, myosin motors, and TMIE provide the first pieces of evidence in support of a new model of slow adaptation. The new model is more aligned with direct PIP_2_ modulation of the channel complex, which is also seen with other ion channels. Inherent mechanosensitivity of the complex may also have a role in mediating the slow adaptation process. The advent of new experimental techniques and expression systems will continue to elucidate the exquisite regulation of the hair cell MET process and allow continued understanding of slow adaptation and other MET regulation mechanisms.

## Materials and Methods

### Tissue preparation

Animals were euthanized using methods approved by the University of Colorado Institutional Animal Care and Use Committee (IACUC). Organs of Corti were dissected in extracellular solution from postnatal (P) day 4 to 5 *Tmie^R82C/+^* and *Tmie^R82C/R82C^*, P7-P9 wildtype (WT) and *Myo7a-ΔC* mice, P7-P10 *Myo1c^Y61G/Y61G^*, and P6-8 Sprague-Dawley rats and placed in recording chambers. Organs of Corti were isolated from the temporal bone and dissected from the modiolus. The tissue was cut at approximately 180 degrees with the mid-apical region as the center. Reissner’s membrane was separated from the lateral wall. The tectorial membrane was then carefully detached using fine forceps to avoid damaging the hair cells. The dissected tissue was transferred to a recording chamber, where strands of dental floss were positioned on the modiolar side, and one was placed on the lateral wall for stabilization. For vestibular hair cells, utricles were isolated from mouse temporal bones. Otoconia were removed carefully using fine forceps without touching the epithelial surface. Strands of dental floss were positioned on the tissue to immobilize the epithelium. Tissue was viewed using a 100x (1.0 NA, Olympus, or 1.1 NA, Nikon) water-immersion objective with a Phantom Miro 320s or Veo 410L (Vison Research, Wayne, NJ) camera on a Slicescope (Scientifica) with Olympus optics or FN1 (Nikon).

### Solution and pharmacology

Tissue was maintained in a normal extracellular solution containing the following components (in mM): 140 NaCl, 2 KCl, 2 CaCl_2_, 2 MgCl_2_, 10 HEPES, 2 Creatine monohydrate, 2 Na-pyruvate, 2 ascorbic acid, 6 dextrose (pH=7.4, 300-310mOsm), or a low divalent extracellular solution containing the following components (in mM): 143 NaCl, 2 KCl, 1 CaCl₂, 0.5 MgCl₂, 10 HEPES, 2 creatine monohydrate, 2 Na-pyruvate, 2 ascorbic acid, and 6 dextrose (pH 7.4, 300–310 mOsm).

We used a low divalent extracellular solution for *Tmie^R82C/+^* and *Tmie^R82C/R82C^* experiments only in order to augment the small MET currents in homozygous mutant mice^26,34^. In all other experments, a normal extracellular solution was used. Apical perfusion was done using pipette tips with diameter of 150 – 300 µm to provide localized perfusion to the hair bundles. For PAO experiments, a 100 mM stock solution of PAO in DMSO was made and solubilized with the help of a bath sonicator. Stock solution was diluted 1:1000 into normal extracellular solution for a final concentration of 100 µM. DMSO control solution was 0.1% DMSO in normal extracellular solution. The composition of the 0.1 mM BAPTA intracellular solution was as follows (in mM): 125 CsCl, 3.5 MgCl₂, 5 ATP, 5 creatine phosphate, 10 HEPES, 0.1 Cs₄BAPTA, and 3 ascorbic acid, pH 7.2, 280–290 mOsm. The composition of the SO₄²⁻ intracellular solution was as follows (in mM): 60 mM Cs_2_SO_4_, 48.8 CsCl, 3.5 MgCl₂, 5 ATP, 5 creatine phosphate, 10 HEPES, 0.1 cesium BAPTA, and 3 ascorbic acid, pH 7.2, 280–290 mOsm. For PIP_2_ experiments, stock solution of 25 mM or 200 mM were prepared by dissolving PIP_2_ (A85155, Avanti Polar Lipids) in ddH_2_O water. On the day of the experiment, the stock solutions (25 mM or 200 mM) were diluted in either 0.1 mM BAPTA intracellular solution or SO₄²⁻ intracellular solution to prepare working solutions with final concentrations of 25 µM, 100 µM, or 200 µM. In the iontophoresis experiments, we used 100 mM EDTA with 25 mM KCl for the pipette solution.

### Electrophysiological recordings

Whole-cell patch clamp experiments were conducted on the first or second row of outer hair cells or type II extrastriolar vestibular hair cells using an Axon 200B amplifier or a Multiclamp 700B (Molecular Devices). Initially, a patch pipette filled with extracellular solution was used to clean the epithelium, and 2-3 adjacent cells near the target cell were removed. Subsequently, a patch pipette filled with intracellular solution was used to perform the whole-cell patch clamp on the cell. Apical perfusion was continuously maintained until a giga-seal was achieved. After breakthrough of the patch membrane to enter whole cell configuration, a minimum waiting time of 5 minutes allowed for the intracellular solution to equilibrate. A fluid jet, using thin-walled borosilicate pipettes (World Precision Instrument), was then used to deliver step-like force stimuli to the hair bundle. Experiments were performed at 18-22°C. Whole-cell currents were filtered at 10 kHz and sampled at 50 kHz using USB-6356 or USB6366 (National Instruments) controlled by jClamp (SciSoft Company). All experiments used -80 mV holding potential unless otherwise noted and did not account for the liquid junction potential. In the LDM experiment, the MET current was first measured at -80 mV using step-like force stimuli, followed by LDM at +80 mV for approximately 10 seconds. After LDM, MET current was measured again at -80 mV with step-like force stimuli. We adopted the results measured at -80 mV after LDM as noted. Iontophoresis experiments were done with pipette resistances in the range 5-7 MΩ. Iontophoresis retention current was 10 nA and ejection currents of 700-1000 nA was applied in increments of 1 second until I_max_ was near 100 pA.

### Hair-bundle stimulation and motion recording

For most of the electrophysiological recordings, hair bundles were stimulated with a custom three-dimensional printed fluid jet driven by a piezoelectric disc bender (27 mm, 4.6 kHz; Murata Electronics or PUI Audio). Thin-wall borosilicate pipettes were pulled to tip diameters of 5 to 15 µm, filled with extracellular solution, and mounted in the fluid-jet stimulator. The piezoelectric disc bender was driven by waveforms generated using jClamp. Stimulus waveforms were filtered using an eight-pole Bessel filter at 1 kHz (L8L 90PF, Frequency Devices Inc.) and variably attenuated (PA5, Tucker Davis) before being sent to a high-voltage/high-current Crawford amplifier to drive the piezoelectric disc bender. During stimulations, videos were taken of the hair-bundle motion with high-speed imaging at 10,000 frames per second using a Phantom Miro 320s or Veo 410L camera (Vision Research) when illuminated with a TLED+ with an LED centered at 530 nm (Sutter Instruments). Videos were saved for each stimulation and analyzed offline. Hair bundle displacement was determined as described previously^10,27,67^. Briefly, hair bundle position was extracted using a Gaussian fit to a high-pass–filtered hair-bundle image for a given vertical row of pixels in the image.

### Immunolabeling and Imaging

Organs of Corti were dissected from wildtype rats and mice. PIP_2_ antibody labeling was performed as described previously^12^. Tissue was fixed in 4% paraformaldehyde (diluted from 16% stock solution, Electron Microscopy Sciences) in 0.1M sodium phosphate buffer (diluted from 0.5M sodium phosphate buffer, Thermo scientific, pH 7.2) for 15 min. After fixation, samples were washed three times with 0.1 M sodium phosphate buffer for 5 min each. The samples were then incubated in 1 % sarkosyl (Bioworld) in PBS for 1 h, followed by three times washes with 0.1 M sodium phosphate buffer for 5 min each. Blocking was performed using 5 mg/ml BSA (Fisher scientific) in TBS (10 mM Tris-HCl, 150 mM NaCl, pH 7,4) for 1 h at room temperature. The samples were then incubated with anti-PIP_2_ monoclonal antibody (1:200 dilution, clone 2C11; Invitrogen) in blocking solution at 4 ℃ overnight. After incubation, the samples were washed three times with blocking solution for 5 min each. Secondary antibodies (1:200 Goat anti-mouse IgM 488; A21042 Invitrogen) in blocking solution were applied for 1 h in the dark. The samples were subsequently washed three times with blocking solution for 5 min each. Alexa Fluor 546-conjugated phalloidin (1:500; Invitrogen) in TBS was applied for 20 min in the dark, followed by three washes with TBS for 5 min each. Tissue was mounted using Prolong Glass Antifade Mountant (Invitrogen). Slides were imaged using a Yokogawa CSU10 mounted on a Nikon FN1 microscope with a 60x oil immersion objective (1.4 NA) and an Aries 16 sCMOS camera (Tucsen Photonics) controlled by µManager^68^.

### Protein Expression and Purification

Full-length murine TMC1 (MmTMC1 FL) was expressed and purified as previously described^69^. Briefly, plasmid containing codon-optimized full-length MmTMC1 (1-757) with an N-terminal hexahistidine-eGFP tag was cloned in the BamHI/NotI sites of the pFastBac1 vector and synthesized by Genscript. Bacmids were generated by transforming plasmids into DH10Bac (ThermoFisher #10361012) cells. *Spodoptera frugiperda* (Sf9) cells were transfected using Cellfectin II reagent (ThermoFisher # 10362100) to produce baculovirus particles, which were amplified through several rounds of infection to yield P2 baculovirus.

To purify MmTMC1 FL, Sf9 cells were infected with P2 baculovirus for 1 day at 27 °C followed by 1 day at 20 °C. Protein expression was monitored by GFP signal. Cell pellets were flash-frozen in liquid nitrogen and stored at -80 °C until needed. Cell pellets were resuspended in 25 mL of solubilization buffer (50 mM Tris-HCl pH8; 150 mM NaCl; 20 mM imidazole; 1 mM beta-mercaptoethanol; and 1 Roche complete EDTA-free protease inhibitor cocktail tablet (Millipore sigma #11836170001)), lysed by probe sonication, and solubilized at 4 °C overnight in n-dodecyl-ß-D-maltoside and cholesteryl hemisuccinate (DDM/CHS from Anatrace #D310 and #CH210, 2/0.4 % w/v). The supernatant was isolated by 1 h ultracentrifugation at 40K rpm at 4 °C, then incubated with 3 mL of Ni-NTA agarose beads (Qiagen #30210) for 3-5 h at 4 °C. Beads were washed once (20 mM Tris-HCl Ph8; 300 mM NaCl; 0.05/0.01% DDM/CHS; 1 mM BME; 20 mM imidazole), then eluted (20 mM Tris-HCl pH8; 300 mM NaCl; 0.05/0.01% DDM/CHS; 1 mM BME; 500 mM imidazole). Eluates were further purified by size exclusion chromatography on a Superose 6 increase 10/300 GL column (Cytiva #29091596) in buffer (20 mM Tris-HCl pH8; 300 mM NaCl; 0.05/0.01% DDM/CHS; 2 mM DTT) and stored at -80 °C until lipid strip experiments.

Protein concentration was measured as absorbance at 280 nm using a spectrophotometer (DeNovix DS-11+), and protein purity was verified using SDS-PAGE gels in NuPAGE Bis-Tris 4-12% gels (ThermoFisher #NP0322), MOPS running buffer (ThermoFisher #NP000102), and PageRuler Plus protein ladder (ThermoFisher #26619).

### Lipid strip assays

Lipid strip assays were performed using the PIP lipid strips (Echelon Biosciences #P-6001). Each strip contains 15 spots with 100 pmol of unique lipids on nitrocellulose membranes and a blank control lacking lipids. GST-tagged PLC-δ1 PH domain protein (PI(4,5)P2-Grip, Echelon Biosciences #G-4501) was used as a positive control for PI(4,5)P2 binding. The protein of interest (4 µg of MmTMC1 or 1 µg of PIP grip) was spotted onto the membrane below the blank control to aid in quantification. Membranes were blocked with lipid strip buffer (PBS with 0.01% Triton X-100 containing 3 % fatty acid-free Bovine Serum Albumin (BSA) (Millipore, #A7030)) for 30 min at room temperature (RT) under gentle orbital shaking. Proteins were added to the membrane at a concentration of 1 µg/mL (PIP grip) or 25 µg/mL (MmTMC1 FL) in lipid strip buffer for 1 h at RT, then washed three times (5 min each) in PBS with 0.01 % Triton X-100 (PBS-T). Primary antibodies were diluted in lipid strip buffer (1:500-1:1000) and incubated with membranes for 1 h at RT. Following another round of washes, membranes were incubated with secondary antibodies diluted in lipid strip buffer (1:2000) and incubated for 1 h at RT. After a final round of washes, Radiance Plus and Radiance Peroxide (Azure Biosystems #AC2103) were combined 1:1 and used for chemiluminescent detection. Antibodies used for MmTMC1 FL were anti-His (primary, ThermoFisher #MA1-21315) and anti-Mouse (secondary, Cytiva #NA931); for the PIP grip, they were anti-GST (primary, ThermoFisher #A-5A00) and anti-Rabbit (secondary, Cytiva #NA9340). An Azure 400 imaging system with automatic acquisition settings was used to image the lipids strips. FIJI was used for quantification of lipid strips. In brief, the average signal from the round ROI around the blank control spot was subtracted from all other spots. Then, intensities were converted to µg of protein using the protein of interest spot. Any negative signal was interpreted as no protein bound (0 µg).

### Methods for Computational Electrophysiology

Microsecond-scale all-atom molecular dynamics (MD) simulations were performed to study how PIP_2_ engages key residues in TMC1 and TMIE and how these interactions evolve under an applied transmembrane potential (-500 mV). The starting structural model was the *Mus musculus* MET channel reported by De-la-Torre *et al.*^43^, assembled as a 2:2:2 complex comprising two TMC1 subunits (TMC1-A and TMC1-B), two TMIE subunits (TMIE-C and TMIE-D), and two CIB2 subunits (CIB2-E and CIB2-F). A membrane environment was generated from this assembly by embedding the MET complex into a POPC bilayer. PIP_2_ was incorporated at 2% of the inner-leaflet phospholipid composition (corresponding to five PIP_2_ molecules), consistent with physiological estimates of PIP_2_ abundance in the plasma membrane^70,71^ and with established plasma-membrane simulation models using comparable concentrations^72^. The system was then neutralized and brought to physiological ionic strength with KCl (0.150 M), and four Ca^2+^ ions were included, two per each TMC1 protomer pore, to populate the pore region during the simulations.

Simulations were performed with GROMACS (v2023.5) using the CHARMM36 force field with CMAP corrections and periodic boundary conditions^73^. Bonds involving hydrogen atoms were constrained with LINCS, enabling a 2-fs integration time step. Long-range electrostatics were treated with Particle Mesh Ewald (PME), and non-bonded interactions were computed with a 1.2-nm real-space cutoff using a Verlet neighbor-list scheme. Temperature was maintained at 310 K using the stochastic velocity-rescaling thermostat with separate coupling groups for protein, membrane, and solvent/ions. Pressure coupling followed a standard membrane protocol, using semi-isotropic Berendsen coupling during equilibration and Parrinello-Rahman coupling during production.

The system followed a six-stage protocol comprising energy minimization, four equilibration steps, and a production stage. After steepest-descent minimization, equilibration was performed in the NPT ensemble for a total of 13 ns (5 ns + 3 ns + 3 ns + 2 ns). During equilibration, harmonic positional restraints were applied to stabilize the MET assembly while allowing solvent and lipids to relax. Protein restraints were applied in a region-specific manner using three force-constant tiers (4200, 2100, and 500 kJ × mol^-1^ × nm^-2^), and the pore sector was additionally restrained with a weaker constant (250 kJ × mol⁻¹ × nm⁻²) to preserve the conductive architecture during early relaxation. Lipid restraints were gradually reduced across equilibration (2000 → 1500 → 1000 → 500 kJ × mol⁻¹ × nm⁻²) and removed for production.

Production simulations were performed under a uniform electric field applied normal to the membrane, corresponding to an effective transmembrane potential of approximately -500 mV. Hair cells sit at a resting potential of roughly −40 to −60 mV ^74^; this gradient drives K^+^ and Ca^2+^ through the open MET channel and. In all-atom MD simulations, physiological transmembrane voltages (−40 to −60 mV) yield insufficient ion permeation events within accessible simulation timescales (ns–μs). Applied electric fields of ±500 mV are routinely used to accelerate ion flux while preserving the directionality and qualitative character of ion conduction ^75–77^.

The production phase was conducted in the NPT ensemble (310 K; Parrinello-Rahman pressure coupling), and the restraint scheme was adjusted to enable pore dynamics: the pore region was fully released, while selected extra-pore regions retained harmonic restraints (Supplemental Figure S8). This strategy was used to maintain global structural stability over microsecond timescales and prevent non-physical deformations that could affect local analyses. Six independent replicas were simulated with distinct initial velocities, five up to 1.5 μs and one up to 2.5 μs. In total we simulated 10 μs of the MET complex interacting with the membrane.

To characterize PIP_2_-protein coupling, trajectories were analyzed for persistent polar and ionic contacts between PIP_2_ headgroups and MET residues. Hydrogen bonds networks were segmented into 10-ns bins to compute binned occupancies and interactions across replicas. For each arginine residue studied, the distance between the guanidinium carbon (Cζ) and the phosphorus atoms of the PIP_2_ P4 and P5 phosphate groups was extracted from the MD simulation trajectories. Because the H-bond contact counts (recorded in 10 ns bins) are reported per residue without distinguishing which of the two phosphate groups were engaged, the distance metric used for comparison with bond counts was defined as the minimum of the instantaneous Cζ–P4 and Cζ–P5 distances, d(t) = min[d_P4_(t), d_P5_(t)]. This choice reflects the fact that bond formation at any given instant is governed by whichever phosphate group is physically closest; the distance to the non-interacting phosphate does not contribute mechanistically to that contact. For each 10 ns bin, the mean of d(t) across all frames within the bin was computed and plotted alongside the corresponding H-bond count for that bin, allowing the time-resolved relationship between proximity and contact formation to be visualized for each analyzed residue.

### Analysis and Statistics

Adaptation time constants were determined for cells that exhibited greater than 20% adaptation magnitude, since smaller adaptation magnitudes often led to poor fits. Current traces were fit with an exponential decay y = y_0_ + A_1_\**e*^-(x–x0)/τ1^. Activation curves were fit with a double Boltzmann equation as described previously^10,22,27^. N represents the number of recorded cells with the number of animals indicated in the figure legend. Data are shown as individual datapoints with summary plots showing mean ± standard deviation for each group. Statistical analysis used Student’s two-tailed *t* tests from MATLAB or Excel. Paired *t* tests were used when comparing data points in the same cell, and unpaired-unequal variance tests analyses were used to compare data between cell populations. Significance (*p* values) are as follows: \**p* < 0.05, \*\**p* < 0.01, \*\*\**p* < 0.001, \*\*\*\**p* < 0.0001. All figures were made using Adobe illustrator.

## Acknowledgements

The authors would like to thank Peter Barr-Gillespie for providing the *Myo1c^Y61G/Y61G^* mice and Ulrich Mueller for providing the *Tmie^R82C/R82C^* mice. This work was funded by NIH-NIDCD R01 DC016868 to AWP, NIH-NIDCD R21 DC019701 to GAC, NIH-NIDCD R01 DC014254 to JBS, and National Research Foundation of Korea (NRF) grant from the Korea government (MSIT) No. RS-2024-00406555 and No. 2022R1A2C3003700 to UKK. GO-O thanks the financial support from the ANID through the FONDECYT de Postdoctorado Project No. 3250793. DR thanks the financial support from the ANID through the FOVI No. 240021. YCP and AB were supported by the Division of Intramural Research of the National Institute on Deafness and Other Communication Disorders (NIDCD DIR DC000096 to AB). The contributions of the NIH authors were made as part of their official duties as NIH federal employees, are in compliance with agency policy requirements, and are considered works of the US government. However, the findings and conclusions presented in this paper are those of the author(s) and do not necessarily reflect the views of the NIH or the US Department of Health and Human Services.

## Author Contributions

GAC and AWP conceptualized and designed the experiments. GAC, SJ, YK, YCP, and AWP conducted experiments and analyzed data. GO-O, CM-G, and DR conducted the computational experiments and analyzed the data. SL and JS generated resources and contributed reagents. AB, UK, JS, DR, and AWP provided funding for the work. GAC, YK, SJ, GO-O, and AWP drafted and revised the manuscript. CM-G, YCP, AB, DR, UK, and JS revised the manuscript.

