## Supplemental Figures for "PIP_2_-TMIE Interactions Drive Mammalian Hair Cell Slow Adaptation Independently of Myosin Motors"

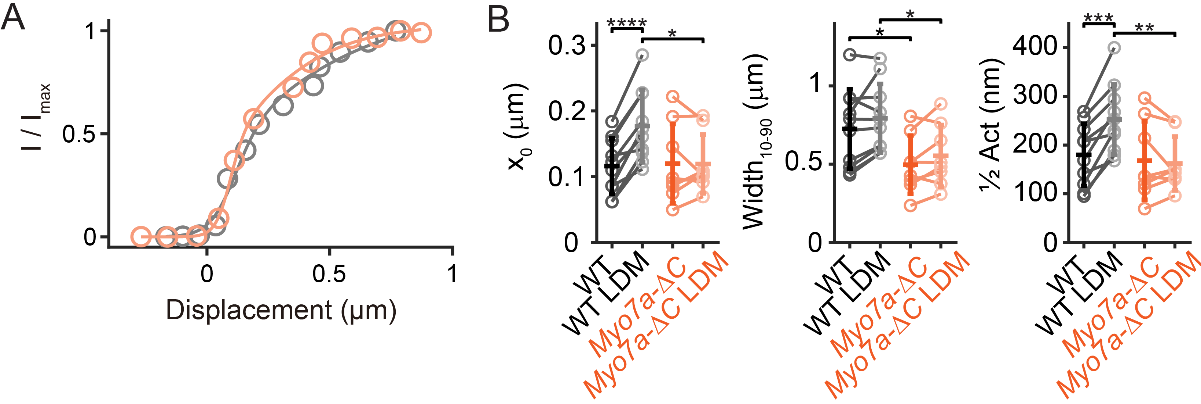

**Supplementary Figure S1**. (A) Activation curves for the cells in Figure 1A and (B) a summary of activation curve parameters for all cells. Data from before LDM and after LDM are connected with lines for WT and *Myo7a-ΔC* hair cells. **p* < 0.05, ***p* < 0.01, ****p* < 0.001, *****p* < 0.0001

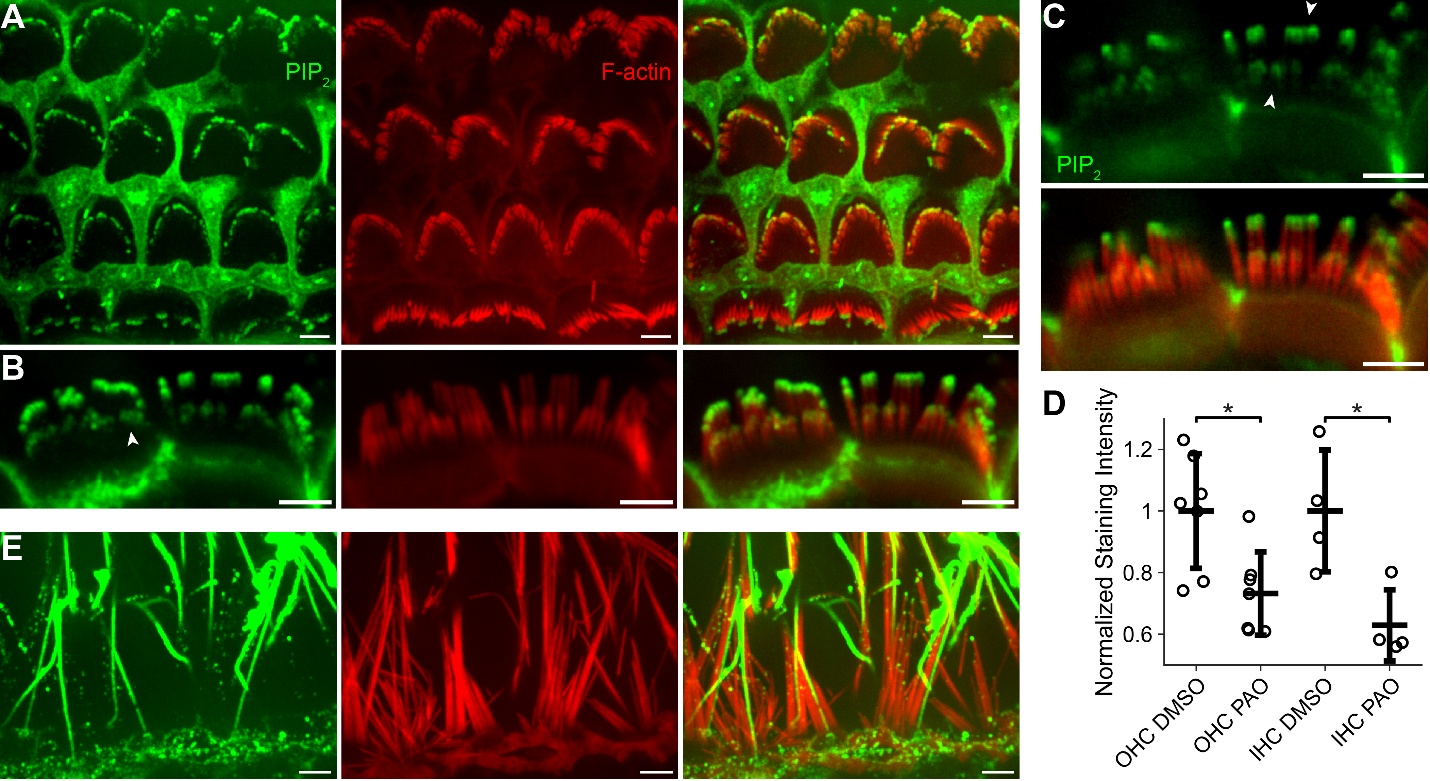

**Supplementary Figure S2**. (A) PIP_2_ labeling in P8 Rat cochlear hair cell stereocilia showing tip labeling in multiple rows of stereocilia. (B) Region of IHCs showing PIP_2_ labeling in all stereocilia tips including the shorter stereocilia (arrowheads). (C) Single optical section of the same hair bundles in (B) showing the cap like labeling pattern (arrowheads). (D) PIP_2_ labeling intensity in stereocilia in control (DMSO treated) and PAO treated OHCs and IHCs show reduced labeling after PAO treatment. (E) PIP_2_ labeling in the vestibular ampulla hair cells also demonstrate stereocilia tip labeling. What appears to be kinocilia are also strongly labeled in vestibular epithelia. Scale bars = 5 µm.

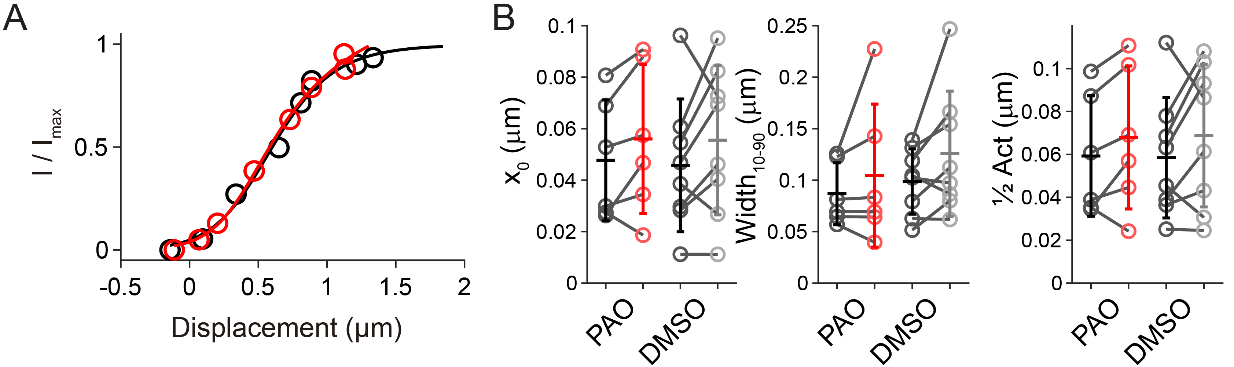

**Supplementary Figure S3**. (A) Activation curves for the cell in Figure 2A and (B) a summary of activation curve parameters for all cells. No significant differences were observed in the activation curve.

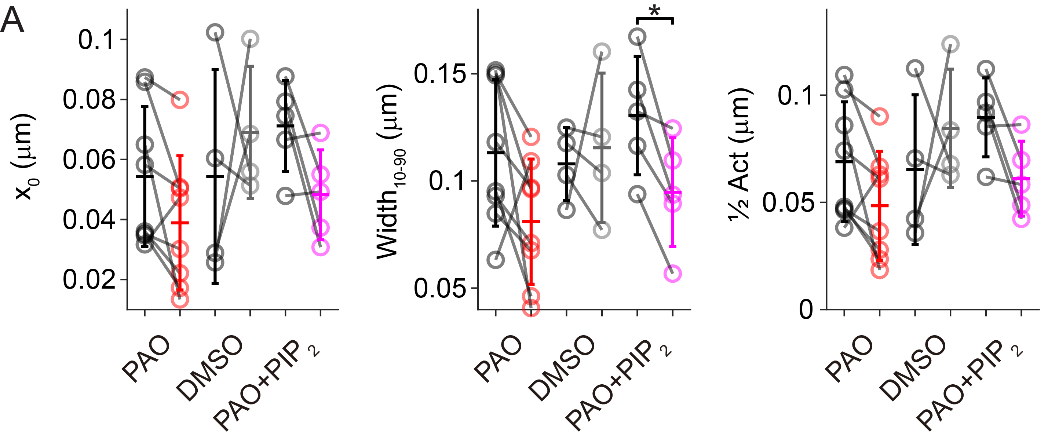

**Supplementary Figure S4**. (A) Activation curve parameters for the 1^st^ activation curve in the multi-pulse protocol before the adapting force step for Figure 3. No significant differences were observed in the activation curve before and after PAO treatment, but there was a significant difference after PAO treatment with PIP_2_ in the pipette for the width of the activation curve. **p* < 0.05

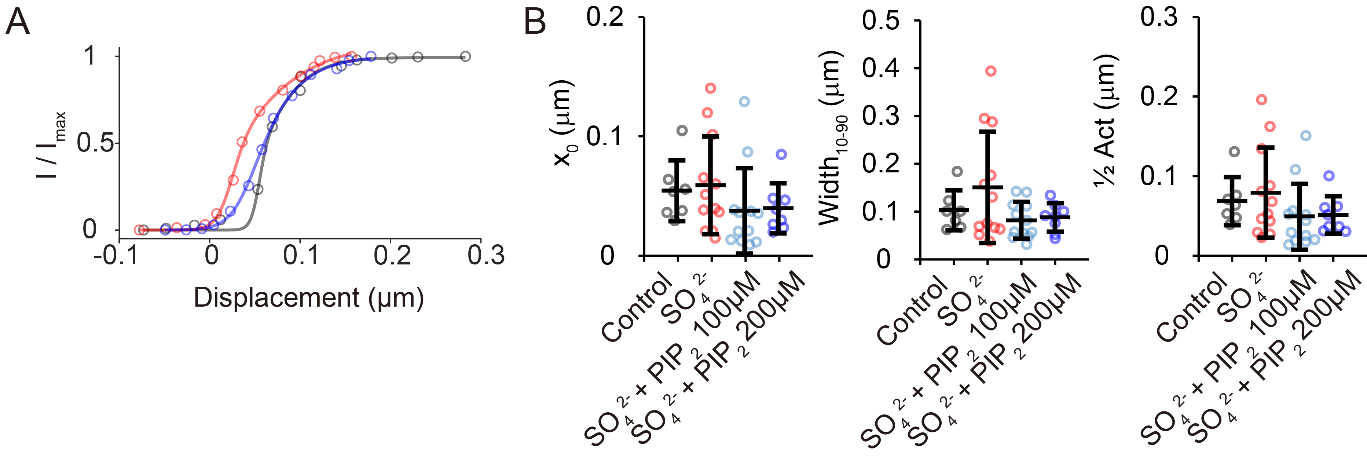

**Supplementary Figure S5**. (A) Activation curves for the cells in Figure 5A-C and (B) a summary of activation curve parameters for all cells. No significant differences were observed in the activation curve.

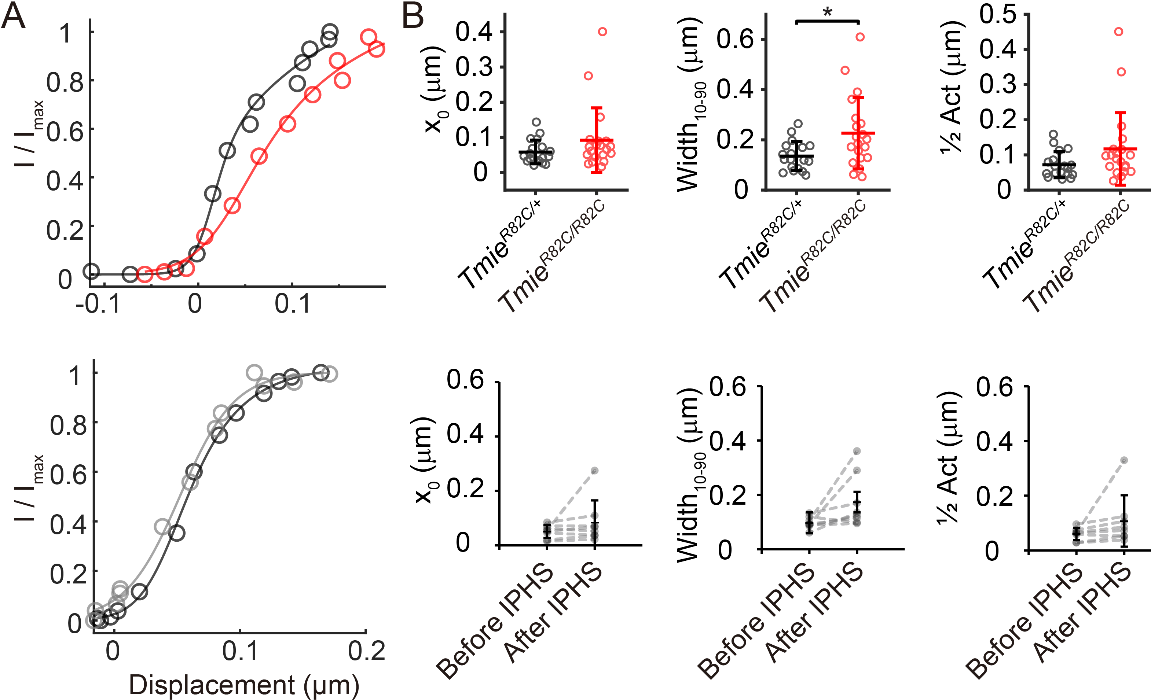

**Supplementary Figure S6**. (A) Activation curves for the cells in Figure 6A, B and (B) a summary of activation curve parameters for all cells. Activation curve width is significantly wider in *Tmie^R82C/R82C^* hair cells. **p* < 0.05.

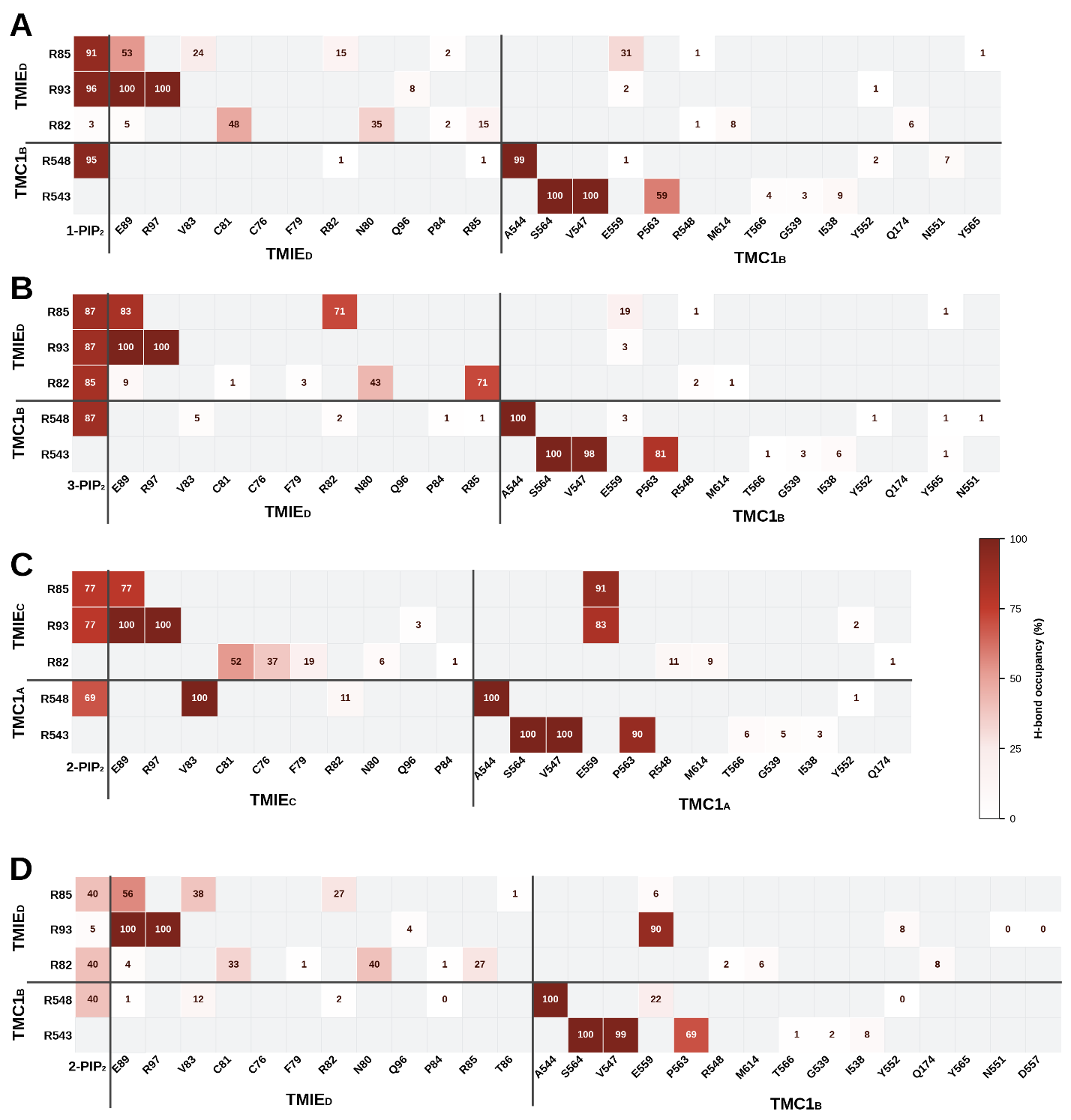

**Supplementary Figure S7.** Hydrogen-bond occupancy profiles of key arginine residues at the TMIE/TMC1/PIP_2_ interface across three independent molecular dynamics replicas. Each panel shows a contact-occupancy matrix in which rows correspond to the arginine residues in the region of interest (R85, R93, and R82 in TMIE; R543 and R548 in TMC1) and columns correspond to their respective interaction partners, grouped by molecular entity. The color intensity of each cell encodes the H-bond occupancy (%), defined as the percentage of 10 ns time bins in which at least one hydrogen bond was detected between the row residue and the column partner, out of the total number of bins spanning the full simulation trajectory. The numerical value inside each colored cell reports this percentage directly; empty grey cells indicate pairs for which no hydrogen bond was detected within the top-10 most-contacted partners for that residue in that replica. The color scale ranges from white (0%) to dark red (100%). (A, C) 1.5 µs replica simulated with two PIP_2_ molecules in both binding sites of the Met complex. (B) 1.5 µs replica simulated one PIP_2_ molecule at one binding pocket. (D) 2.5 µs replica simulated with one PIP_2_ molecule at the binding pocket. Panels A, B, C, and D correspond to Figures 7D, 7E, 7F, and 7G of the main text, respectively. Panels A–D collectively demonstrate that the hydrogen-bond network engaging R85, R93, and R548 with PIP_2_ and with neighboring TMIE and TMC1 residues is a reproducible and persistent feature across independent simulation conditions and trajectory lengths. The intra-TMC1 contacts of R543 with S564, V547, and P563 reach near-complete occupancy (≥90%) in all four replicas, indicating a structurally stable hydrogen-bond network within the TMC1 subunit that is independent of PIP_2_ composition. All simulations were performed using the CHARMM36 force field with GROMACS.

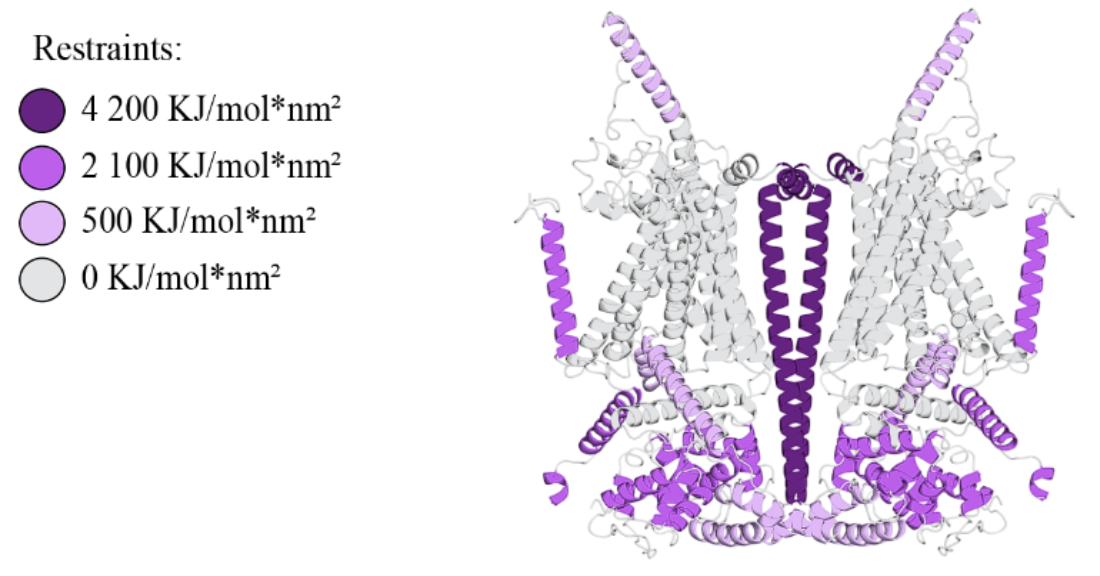

**Supplementary Figure S8**. Positional restraint scheme used during equilibration and production MD simulations of the MET complex. Protein regions were restrained with harmonic positional potentials using the force constants indicated in the legend. During equilibration, the pore sector (grey) was additionally restrained with k = 250 kJ·mol^-1^nm^-2^ to preserve the conductive architecture while the membrane and solvent relaxed.

**Supplementary Video 1.** Video of 1.5 µs MD simulation corresponding to Figure 7F showing the interactions of the PIP2 headgroup with R85 and R93 of TMIE and R548 of TMC1.
